# Analysis of Wilms’ tumor protein 1 specific TCR repertoire in AML patients uncovers higher diversity in patients in remission than in relapsed

**DOI:** 10.1101/2023.11.26.568717

**Authors:** Sofie Gielis, Donovan Flumens, Sanne van der Heijden, Maarten Versteven, Hans De Reu, Esther Bartholomeus, Jolien Schippers, Diana Campillo-Davo, Zwi N. Berneman, Sébastien Anguille, Evelien Smits, Benson Ogunjimi, Eva Lion, Kris Laukens, Pieter Meysman

## Abstract

The Wilms’ tumor protein 1 (WT1) is a well-known and prioritized tumor-associated antigen expressed in numerous solid and blood tumors. Its abundance and immunogenicity have led to the development of different WT1-specific immune therapies. The driving player in these therapies, the WT1-specific T-cell receptor (TCR) repertoire, has received much less attention. Importantly, T cells with high affinity against the WT1 self-antigen are normally eliminated after negative selection in the thymus and are thus rare in peripheral blood. Here, we developed computational models for the robust and fast identification of WT1-specific TCRs from TCR repertoire data. To this end, WT1_37-45_ (WT1-37) and WT1_126-134_ (WT1-126)-specific T cells were isolated from WT1 peptide-stimulated blood of healthy individuals. The TCR repertoire from these WT1-specific T cells was sequenced and used to train a pattern recognition model for the identification of WT1-specific TCR patterns for the WT1-37 or WT1-126 epitopes. The resulting computational models were applied on an independent published dataset from acute myeloid leukemia (AML) patients, treated with hematopoietic stem cell transplantation, to track WT1-specific TCRs *in silico*. Several WT1-specific TCRs were found in AML patients. Subsequent clustering analysis of all repertoires indicated the presence of more diverse TCR patterns within the WT1-specific TCR repertoires of AML patients in complete remission in contrast to relapsing patients. We demonstrate the possibility of tracking WT1-37 and WT1-126-specific TCRs directly from TCR repertoire data using computational methods, eliminating the need for additional blood samples and experiments for the two studied WT1 epitopes.

## Introduction

Central in the development of new immunotherapies for patients with acute myeloid leukemia (AML) lies the specific targeting of AML-associated antigens [1], [2]. A specific interest goes to Wilms’ tumor protein 1 (WT1) which has an acknowledged role as a tumor oncogene in a variety of malignancies, including AML [2], [3]. As such, WT1 has been identified as a nearly universal tumor-associated antigen (TAA) overexpressed in numerous solid and hematological cancers [4]–[7], and has been listed as the most interesting cancer antigen for immune therapies [8]. However, as a self-antigen also expressed in healthy tissues, T-cell clones of high affinity are usually eliminated after negative selection in the thymus. Therefore, the frequency of high-affinity T-cell receptors (TCRs) towards WT1-epitopes in circulating T cells is low. It is hypothesized that WT1-targeted immunotherapies might increase the frequency of anti-AML T cells activity resulting in higher survival rates. Many of these therapies are based on activation of WT1-specific T cells targeting cancer cells overexpressing the WT1 antigen. Hence, the success of these immune therapies relies on recognition of the WT1 antigen by the TCR of these WT1-specific T cells. Despite the large interest in WT1, the WT1-specific TCR repertoire in AML patients has not been fully investigated so far.

TCRs are heterodimeric membrane proteins consisting most often of an alfa and a beta chain (αβ TCRs), each containing three hypervariable domains called complementary determining region 1 (CDR1), CDR2, and CDR3. The CDRs are in contact with the peptide and/or the major histocompatibility complex (MHC) molecule. Especially the CDR3 alpha and beta regions are important in identifying its epitope partner as these largely interact with the epitope surface [9], [10]. Due to the high diversity of these CDR3 sequences in each TCR repertoire, the immune system is able to recognize a broad spectrum of peptides. This TCR diversity is achieved through TCR gene rearrangement, a process in which discrete segments composing the TCR genes (V and J in the alfa chain, and V, D and J in the beta chain) randomly join by somatic recombination. The introduction of short insertions and deletions at the connections between rearranged genes further increases TCR diversity [11].

TCR sequencing on blood samples and biopsies facilitates the study of the composition of this repertoire. In this sense, to analyze the TCR repertoire of a sample, both bulk and single-cell sequencing are used. While the former strategy is less expensive per T cell, the latter allows the recovery of paired TCR alpha-beta chains. In addition, antigen-specific T cells can be isolated by sorting tetramer-positive T cells specific for certain peptide-MHC complexes [12] with or without concomitant selection of T cells that upregulate the expression of activation markers such as CD137 [13], [14]. T-cell sorting is followed by TCR sequencing, aiding the study of antigen-specific TCR repertoires. Due to the increasing availability of public epitope-specific TCR data, new methods were designed to circumvent these laborious experiments by focusing on the computational identification of epitope-specific TCRs. This progress was possible since TCRs with similar CDR3 beta sequences are known to often recognize the same epitope [15]. Thus, previously identified epitope-specific TCRs can be used to identify common patterns that underlie the recognition of epitopes by TCRs. This is the basis of tools such as TCRex [16], which use machine learning methods to extract epitope-specific recognition patterns in the TCR CDR3 sequence to predict the epitope specificity for new TCRs. In addition, the finding of shared similar patterns in the sequences of epitope-specific TCRs has led to the development of clustering methods such as clusTCR [17] and ALICE [18]. These allow the identification of potentially epitope-specific clusters by grouping TCRs based on their CDR3 sequence.

In this study, we applied recently developed computational techniques to study the TCR repertoire for two human leukocyte antigen (HLA)-A*02:01-restricted WT1-derived peptides: WT1_37-45_ and WT1_126-134_, hereafter referred to as WT1-37 and WT1-126, respectively [19], [20]. The objective was to train machine learning methods to monitor WT1-specific T-cells in the repertoires of patients with AML.

## Materials and methods

A summary of the general workflow is demonstrated in Figure 1. A full description of the materials and methods is present in **Supplemental material S1**.

**Figure 1:**
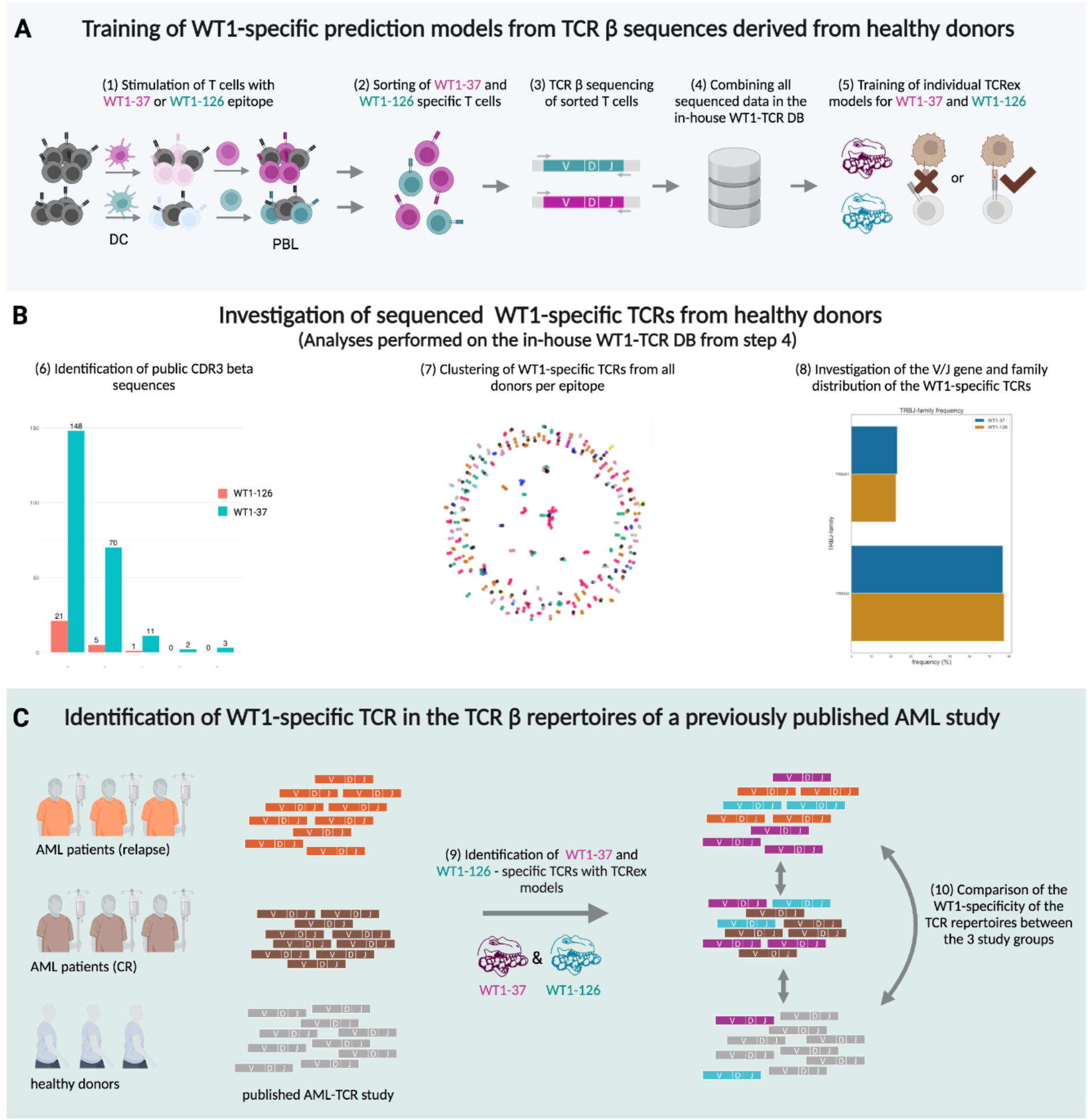
Graphical overview of the different methodologies and data sets. **(A) Collection of WT1-specific T cells from healthy donors for prediction model training.** (1) WT1-specific CD8^+^ T cells were expanded from HLA-A*02:01-positive healthy donor buffy coats by two consecutive *in vitro* stimulations (IVS) [20] and (2) subsequently sorted with epitope-specific MHC class I tetramers. Following TCR β sequencing of the sorted T cells (3), the resulting WT1-specific TCRs were combined into one database (4), referred to as ‘in-house WT1-TCR DB’ and used to train a WT1-37 and WT1-126 TCRex model (5). **(B) In-depth analysis of the TCR β sequences derived from the *in vitro* stimulated T cells as described in (A)**. The presence of patterns shared between donors was assessed by (6) identification of public TCRs and (7) clustering of all WT1-specific TCRs for each epitope separately. (8) V/J gene distribution was also assessed in relation to an independent background data set. **(C) Identification of WT1-specific TCRs in the TCR β repertoires of AML patients and healthy donors.** Usability of the trained TCRex models from step (4) was assessed using an independent published data set [21], containing TCR β repertoires from three healthy donors, three AML patients with relapse and three AML patients showing clinical response following hematopoietic stem cell transplantation (HSCT). Using the trained TCRex models, WT1-37-specific (pink) and WT1-126-specific (blue) TCRs were identified in the repertoires of the healthy donors and AML patients (9) and compared between groups (10). Figure created with biorender.com.

## Results

### Establishment of a WT1-TCR database from expanded WT1 epitope-specific primary human CD8^+^ T cells

WT1-37-reactive and WT1-126-reactive CD8^+^ T-cell clones were successfully expanded using buffy coat preparations by means of two consecutive IVS. After the first IVS with autologous peptide-pulsed monocyte-derived DCs, 0.29 ± 0.23% (mean ± SD) of viable WT1-37-specific (**Figure 2A**) and 0.08 ± 0.11% of WT1-126-specific CD8^+^ T cells (**Figure 2B**) were detected with WT1-37 or WT1-126 HLA-A*02:01 tetramers (WT1-37/HLA-A2 and WT1-126/HLA-A2, respectively). After the second IVS with irradiated autologous peptide-pulsed CD14^-^CD8^-^ peripheral blood lymphocytes, both WT1-37-specific and WT1-126-specific T cells significantly increased to 8.03 ± 7.66% (p=0,0005; **Figure 2A**) and to 1.13 ± 1.35% (p=0,0469; **Figure 2B**), reflecting a mean 28-fold and 14-fold increase in the frequency of WT1-37-specific and WT1-126-specific CD8^+^ T cells, respectively, between the first and the second IVS. Next, the TCR β chains of WT1-37/HLA-A2 and WT1-126/HLA-A2 tetramer-sorted T cells were sequenced. The full list of TCR sequences can be found in **Supplemental material S2**. An overview of the number of raw reads and the number of TCR clonotypes identified by MiXCR is given in **Supplemental material S3**. In total, 1262 and 101 unique TCR β sequences were derived from 12 healthy donors for peptide WT1-37 and seven healthy donors for peptide WT1-126, respectively.

**Figure 2:**
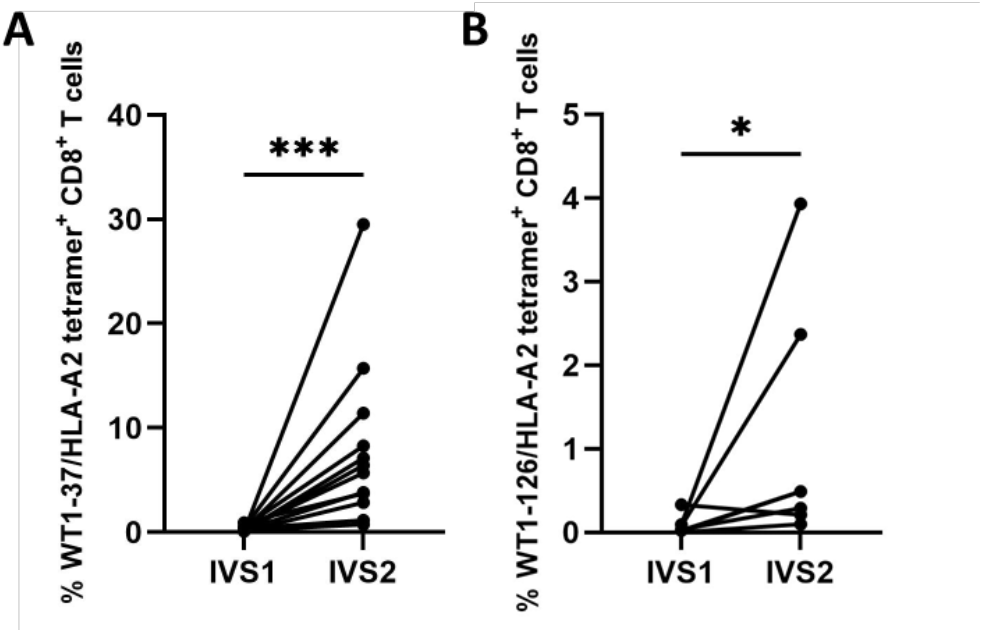
Percentage of expanded WT1-37/HLA-A2 and WT1-126/HLA-A2 tetramer-positive WT1-epitope specific CD8^+^ T cells for subsequent fluorescence-activated cell sorting. Primary peripheral blood CD8^+^ T cells from healthy donors were significantly expanded with **(A)** WT1-37 peptide (n=12, p=0.0005) or **(B)** WT1-126 peptide (n=7, p=0.0469), in two rounds of IVS (Wilcoxon t-test). Viable WT1-37/HLA-A2 and WT1-126/HLA-A2 tetramer-positive CD8^+^ T cells were sorted for RNA extraction and subsequent bulk sequencing. Abbreviations: IVS, *in vitro* stimulation; WT1, Wilms’ tumor protein 1. *, p ≤ 0.05; ***, p ≤ 0.001.

### WT1-specific TCR sequences from different healthy repertoires cluster together

In addition to public TCRs that share identical TCR β sequences (**Supplemental material S4**), it is expected to find similar TCR β sequences that only differ in a small number of their amino acids across the WT1-specific repertoires of healthy donors. To evaluate the level of TCR similarity, all WT1-specific CDR3 beta sequences were clustered using clusTCR [17] (**Figure 3**). **Figure 3** depicts clusters for the 411 WT1-37-epitope specific CDR3 beta sequences out of the total 1262 extracted sequences (33%). **Figure 3B** shows clusters for the 34 WT1-126-epitope specific CDR3 beta sequences out of the total 101 extracted sequences (34%). While some clusters are unique to a single donor, many clusters contain TCRs from two or more different donors (69/166 clusters for WT1-37 and 12/16 clusters for WT1-126), indicating that WT1-specific TCRs share similarities within their CDR3 beta sequences across donors (**Figure 3C**). With the exception of the conserved cysteine and phenylalanine at the beginning and the end of the CDR3 beta sequences, sequence logos (**Supplemental material S5)** show that every other position can be represented by more than one amino acid. There is thus no strict location where the amino acid sequences differ between CDR3 beta sequences of one cluster. Sequence logos also reveal that the varied amino acids at each position are not restricted to a specific group of amino acids, as exemplified in sequence logo **Figure 3C**, position six contains both hydrophobic (cysteine, C) and hydrophilic (arginine, R) amino acids. An overview of the importance and distributions of the V/J genes is given in **Supplemental materials S6-S7.**

**Figure 3:**
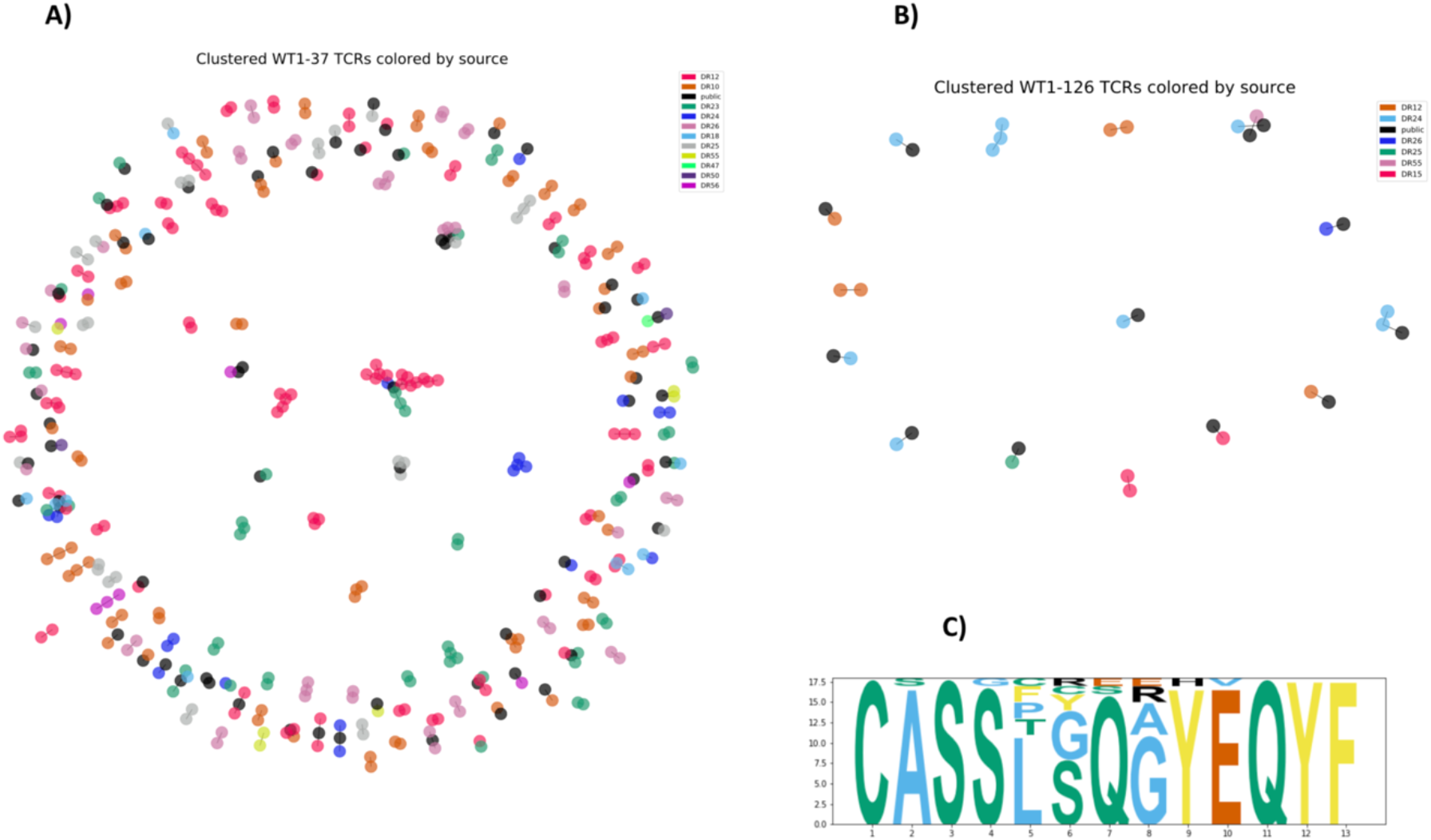
Graphs of the clustered WT1-37 (A) and WT1-126 CDR3 beta sequences (B) colored according to the donors. CDR3 beta sequences shared with more than one donor (i.e., public TCRs) are depicted in black. (**C**) Sequence logo of the largest cluster for WT1-37. In general, the sequence logos summarize the amino acid composition of the CDR3 beta sequences for every cluster (**Supplemental material S5)**. The largest cluster contains 18 CDR3 beta sequences with a length of 13 amino acids. For each of the 13 CDR3 beta positions, the sequence logo summarizes which amino acids appear at these positions in the 18 sequences, while the size of the letters represents the frequency of each amino acid in a particular position. For example, all 18 CDR3 beta sequences contain a cysteine (C) on the first position, while either a serine (S) or a glycine (G) can be present at the fourth position. Since the size of the letter S at position four is much larger than the letter G, most sequences will contain a serine at this position.

### WT1-specific prediction models can be built with good performance

The cluster analysis (**Figure 3**) showed the generalizability of the epitope-specific features in the in-house WT1-TCR DB created using samples from healthy donors as starting material. Next, we analyzed whether the WT1-specific TCR repertoire contained enough epitope-specific information in their CDR3 beta sequences to train WT1-specific prediction models with TCRex. TCRex automatically returns common performance metrics using a 5-fold cross-validation strategy (**Supplemental material S8**) and a receiver operating characteristic (ROC) and precision-recall plot for every trained model (**Figure 4**). For the identification of WT1-specific TCRs in independent data, both models were evaluated. The models showed sufficient performance to be used as a predictor for WT1-specific TCRs.

**Figure 4:**
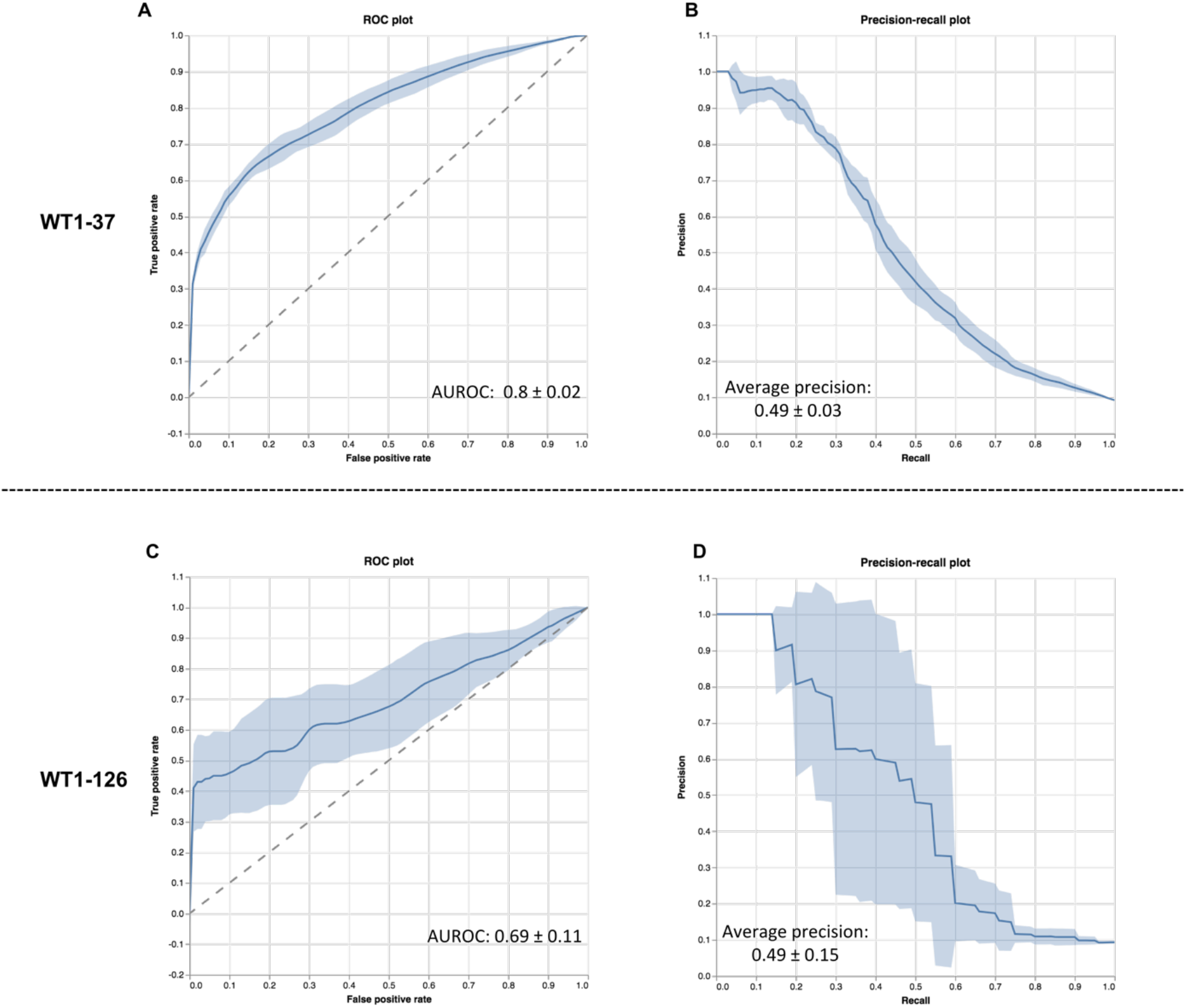
Receiver operating characteristic (ROC) and precision-recall curves for the trained WT1-37 and WT1-126 TCRex models. ROC curves visualize the true positive rate (i.e., fraction of the WT1-specific training data set that is correctly classified as WT1-specific by the model) and false positive rate (i.e., fraction of the background data that is wrongly classified as WT1-specific) for all possible classification thresholds for WT1-37 (**A**) and WT1-126 (**C**) TCRex models. The precision-recall plots depict the precision at different classification thresholds over the true positive rate for WT1-37 (**B**) and WT1-126 (**D**) TCRex models. The precision reflects the proportion of predicted WT1-specific TCRs that is truly WT1-specific and is thus desired to be approximating 1.

### WT1-specific TCRs are identified from independent AML patient TCR repertoire data by the trained prediction models

Two prediction models were trained using TCR beta sequences derived from healthy individuals, one for every WT1 epitope. In order to validate their usability on cancer TCR repertoires, the two models were utilized for the identification of WT1-specific TCRs in cancer patients. For this, we used an independent data set investigating T cells in the bone marrow of AML patients after hematopoietic stem cell transplantation (HSCT) [21]. For both prediction models, WT1-37 and WT1-126 specific TCRs were identified both in one of the healthy individuals and five AML patients, of which two patients were in relapse and three patients in complete remission. AML patients in remission showed the highest frequency of WT1-specific TCRs (nine identified WT1-specific TCRs), compared to healthy individuals and relapsed patients (three and five, respectively), and contained more unique TCRs having a larger TCR repertoire size (**Table 1**). Two of the identified WT1-specific TCR CDR3s were present both in patients in complete remission as well as in relapse (**Supplemental material S9**).

**Table 1:** Identified WT1-specific TCRs in AML patients versus healthy donors.

| Individual | Disease status | Size of the TCR repertoire | Number of WT1-37 TCRs | Frequency of WT1-37 TCRs (%) | Number of WT1-126 TCRs | Frequency of WT1-126 TCRs (%) |
| --- | --- | --- | --- | --- | --- | --- |
| HD1 | Healthy | 295 | 0 | 0 | 0 | 0 |
| HD2 | Healthy | 110 | 0 | 0 | 0 | 0 |
| HD3 | Healthy | 931 | 2 | 0.215 | 1 | 0.107 |
| PT1_CR1 | Complete remission | 3099 | 4 | 0.129 | 0 | 0 |
| PT2_CR2 | Complete remission | 1451 | 3 | 0.207 | 1 | 0.069 |
| PT3_CR3 | Complete remission | 102 | 1 | 0.980 | 0 | 0 |
| PT4_REL1 | Relapse | 503 | 0 | 0 | 0 | 0 |
| PT5_REL2 | Relapse | 592 | 2 | 0.338 | 0 | 0 |
| PT6_REL3 | Relapse | 617 | 2 | 0.324 | 1 | 0.162 |
| For each of the studied individuals, size of the CD4<sup>+</sup>CD8<sup>+</sup> bone marrow TCR repertoire, number and corresponding percentage of identified WT1-TCRs is given for each WT1-epitope individually.<br>Abbreviations: CR, complete remission; HD, healthy donor; PT, patient; REL, relapse. |  |  |  |  |  |  |

### WT1-specific TCR clusters are associated with AML

In case of peptide-specific T-cell expansion, it is expected to find clusters of highly similar TCRs across repertoires that are reacting to the same peptide-MHC complex [18]. In total, 721 of the 7600 unique CDR3 beta sequences (9,5%) were detected in clusters of similar sequences (**Figure 5A**) from TCR repertoires derived from bone marrow samples from an independent data set [21]. 90 clusters combined TCRs from healthy individuals and AML patients, suggesting substantial overlap between these groups. When considering the seven clusters containing at least one predicted WT1-specific TCR (**Table S10 ,** **Figure 5B**), the majority (39 out of 42) were made up only of sequences that were found in AML patients (**Supplemental material S10**), indicating disease-specificity and (predictive) relevance of WT1-specific TCRs in AML. In the three clusters that included TCRs present in healthy samples, the AML-linked TCRs still dominated. Remarkably, TCRs derived from patients in complete remission were spread out over all seven clusters, while TCRs from relapsing patients were concentrated in only three (**Table S10**, **Figure 6**), suggesting that displaying more diverse WT1-specific TCR patterns could be associated with remission (chi-squared test, p=0.0043).

**Figure 5:**
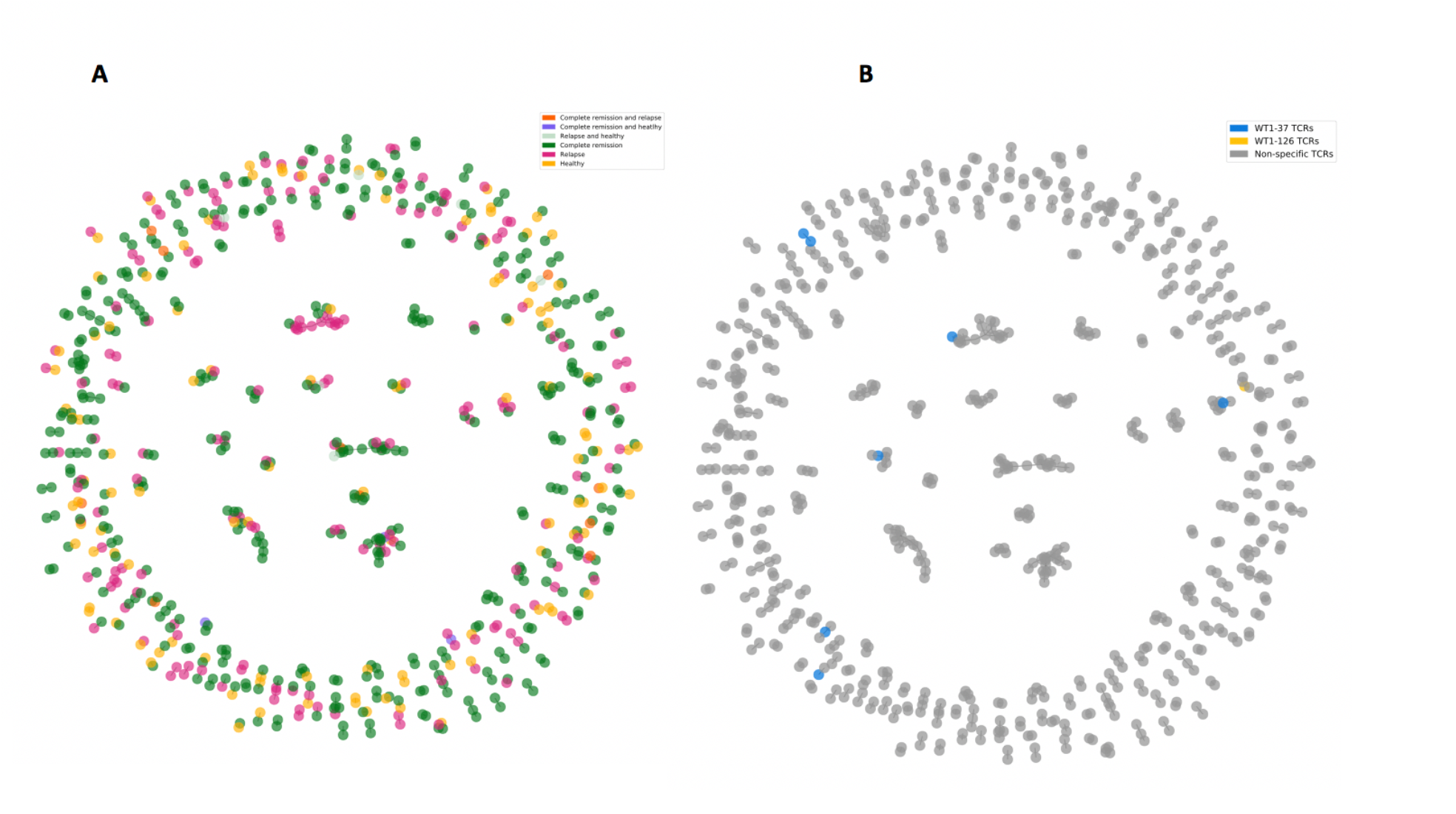
Clustered CDR3 beta repertoires of six AML patients and three healthy individuals from an independent data set. [21]. Clusters are colored by (**A**) response and (**B**) WT1 specificity. Each colored dot represents a single CDR3 beta sequence. Clusters are defined by at least two connected CDR3 beta sequences.

**Figure 6:**
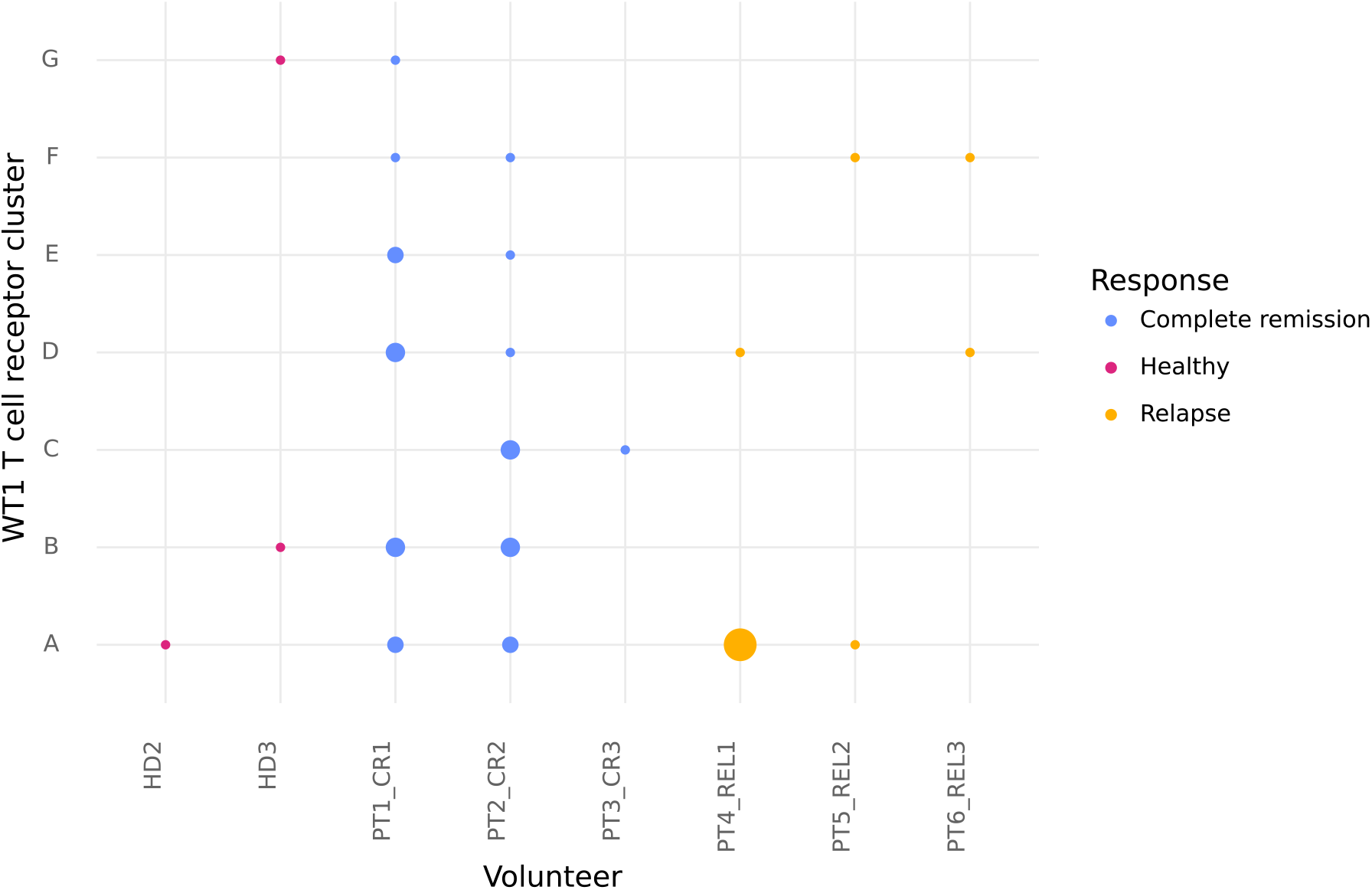
Overview of the WT1-specific TCR clusters. Letters on the y-axis represent a cluster containing at least a single WT1-specific TCR. On the x-axis, every individual containing at least a single WT1-specifc TCR in its repertoire is listed. Out of nine individuals, WT1-specific TCRs were detected in eight individuals. The size of the dots represents the number of TCRs from an individual that are present in that specific cluster, while the color represents the study groups based on disease status: healthy (red), AML patients in complete remission (blue), and relapsed AML patients (yellow). Abbreviations: CR, complete remission; HD, healthy donor; PT, patient; REL, relapse; TCR, T cell receptor; WT1, Wilms’ tumor protein 1.

## Discussion

The greater availability of bulk TCR repertoire sequencing has sparked interest in using the TCR repertoire as a multiplexed diagnostic assay [22]. Specificities against a large collection of epitopes can be assessed from the same TCR repertoire using *in silico* annotation models predicting epitope-TCR pairings. Such annotation models require training on previously identified epitope-specific TCRs for any target epitope, as unseen epitope prediction remains an unsolved problem [23]. However, the limited availability of patient samples for rare malignancies complicates the collection of TCR data for different epitopes. To address this issue, here, we propose a workflow to efficiently isolate WT1-specific TCRs from healthy individuals and train WT1-specificity prediction models. The resulting TCR sequences from *in vitro* expanded T cells from healthy donors showed similarities across donors, which allowed the training and use of WT1-specific models for the identification of WT1-specific TCRs and thus tracking of WT1-specific TCRs in AML patients. By using this approach, we identified more WT1-specific TCRs in the repertoires of the AML patients compared to healthy individuals (12 versus 2). A similar result was found when healthy individuals were vaccinated against Yellow Fever Virus (YFV). Here, the vaccinated repertoires contained more unique YFV-specific TCRs than the pre-vaccinated repertoires [16]. Similar to [24] and [25], we believe that the contact with AML cells stimulates the expansion of WT1-specific TCRs. This is further supported by other studies demonstrating a higher level of WT1-specific T cells in leukemic patients in contrast to healthy individuals [26], [27]. Hence, assessment of WT1-TCR repertoire research as a diagnostic tool for AML might be considered[22].

Along with the annotation of tumor-specific T cells on an individual TCR basis, computational methods are being developed to discover groups of TCRs recognizing the same epitope [17]. These so-called clustering methods were established after the observation that TCRs with similar amino acid sequences frequently show similar epitope specificities [28]–[30]. Big TCR clusters in a repertoire are indicative of a convergent proliferation against a limited set of epitopes and can therefore capture an ongoing or past tumor-specific immune response [18], [31]. During the analysis of potential WT1-specific TCR clusters, we observed that TCRs from relapsed AML patients were more restricted to a limited number of clusters when compared to AML patients in complete remission. Therefore, it is suggested that displaying more diverse WT1-specific patterns, as seen in patients in complete remission, may be associated with a greater protection against relapse. In this sense, it is already known that TCR repertoire diversity plays a role in treatment response. Various studies have shown a link between TCR diversity and response to immune checkpoint inhibitors [32]–[34]. Moreover, a study on DC-based vaccination has demonstrated an increase in the diversity of the melanoma neoantigen-specific TCR repertoire following treatment [35]. In addition, Rezvani et al. demonstrated that the WT1-specific CD8+ T cells of patients with chronic myelogenous leukemia or AML targeted more WT1-epitopes than healthy volunteers [27]. In that line, our group made an association between the levels of WT1-specific CD8+ T cells targeting different WT1-epitopes and the clinical response of AML patients following a WT1-based dendritic cell vaccine [36], while Hoffman et al. discovered that patients responding to donor lymphocyte infusion (DLI) targeted more leukemia associated epitopes than those who did not respond [37]. Taken together, these data underscore the impact of TCR repertoire diversity in targeting different tumor epitopes. In this study, we identified a relation between the diversity of epitope-specific TCR signatures for individual WT1-epitopes, as defined by the presence of TCRs in WT1-specific clusters, and the response to HSCT. This signal is based on the two WT1-epitopes for which we generated data from healthy individuals, and thus represents a fraction of the overall cellular immunity against the WT1 antigen. Further investigation of the full TCR repertoire is warranted, with an extended coverage of WT1-associated epitopes. Hence, the same protocol can be repeated for other WT1-epitopes resulting in additional prediction models expanding the identification of WT1-specific TCRs.

Overall, we demonstrate that the WT1-specific T-cell repertoire of AML patients holds information regarding response to HSCT therapy. Thus, the WT1-specific TCR repertoire has potential as a promising biomarker for response prediction to cancer treatment and could provide invaluable data for the allocation of personalized immunotherapies to patients that would most likely respond to therapy, especially considering the high costs of personalized immunotherapy. In addition to the absolute count of unique WT1-specific TCRs and the diversity of the WT1-specific TCR repertoire, the activity and function of these T cells may also hold information about the different patient groups. Furthermore, no baseline data was available, thus the presence of the WT1-signal repertoires prior to treatment could not be assessed. This information could provide additional insight into the existence of biomarkers in the baseline repertoires and their evolution over time, potentially aiding in patient stratification [38]. In conclusion, we have developed a workflow that starts from a robust antigen-specific primary T-cell expansion platform to collect WT1-specific TCRs from the blood of healthy individuals. The collected TCRs were used successfully to train computational models for the identification of WT1-specific TCRs in independent data sets from AML patients. Our workflow revealed differences in the number of unique WT1-specific TCRs and cluster diversity between patients in complete remission and those experiencing relapse. The described platform is extrapolatable to other cancer antigens, enabling the identification of tumor-specific clonotypes within full TCR repertoires. This can aid the research of biomarkers for cancer immunotherapies.

### Author contributions

S.G. performed research, collected data, analyzed and interpreted data and wrote the original manuscript. D.F. performed research, collected data, analyzed and interpreted data and wrote the original manuscript. S. v.d. H. performed research, reviewed and edited the manuscript. M. V. performed research, collected data and reviewed the manuscript. H. D. R. performed research and reviewed the manuscript. E. B. conducted research, analyzed and interpreted the data and reviewed the manuscript. J. S. performed research. D. C. contributed to the interpretation of the results, reviewed and edited the manuscript. Z. N. B. reviewed and edited the manuscript. S. A. reviewed the manuscript. E. S. reviewed and edited the manuscript and supervised the project. B. O. reviewed and edited the manuscript, helped out in research design, conceived the study and supervised the project. E. L. reviewed and edited the manuscript, helped out in research design, conceived the study and supervised the project. K. L. reviewed and edited the manuscript, helped out in research design and supervised the project. P. M. reviewed and edited the manuscript, helped out in research design, conceived the study and supervised the project.

## Funding

This work was supported in part by research grants of the Special Research Fund of the University of Antwerp (Small Research Grants ID36379, BOF-KP2018 P.M. and ID41605, Concerted Research Action) and the Methusalem financing program of the Flemish Government to the Antwerp University, the Cellular Therapy Fund from the Antwerp University Hospital (UZA) and UZA Foundation (Belgium), iBOF Project “Modulating Immunity and the Microbiome for Effective CRC Immunotherapy” (MIMICRY) and the public utility foundation Stichting ME TO YOU (Belgium). D.F. was supported by the ‘‘Cellular Immunotherapy’’ grant from the Baillet Latour Fund (Belgium), and an Emmanuel van der Schueren fellowship from Kom op tegen Kanker (KotK, Belgium), followed by a DOCPRO PhD grant of the Special Research Fund (Grant MulTplex). S.G. and M.V. were supported by an SB fellowship from the FWO (S.G. grant 1S48819N, M.V. grant 1S24517N). E.B. and S.v.d.H. were supported via a clinical investigator fellowship from the Research Foundation-Flanders (FWO, Belgium) of B.O. (grant 1861219N). D.C.D. and H.D.R. were supported by grant G053518N from the FWO.

### Data availability statement

Data and scripts used for the analysis are available on github: https://github.com/sgielis/WT1_TCR. The trained models are available on https://tcrex.biodatamining.be/. The original MiXCR files from the healthy volunteers are also available on Zenodo (DOI 10.5281/zenodo.8129250)

