## Supplementals for "Analysis of Wilms’ tumor protein 1 specific TCR repertoire in AML patients uncovers higher diversity in patients in remission than in relapsed"

### Supplemental materials

#### S1: Details regarding the materials and methods

##### Expansion and sorting of WT1-specific $CD8^+$ T cells from healthy donors

Buffy coats from healthy anonymous donors were purchased from the Blood Service of the Flemish Red Cross (Mechelen, Belgium) following the approval by the Ethics Committee of the Antwerp University Hospital and the University of Antwerp under reference number 15/19/210. (Informed consent was collected by the Dienst van het Bloed of the Red Cross-Vlaanderen). Whole blood HLA-typing was performed to select for HLA-A*02:01+ donors. Expansion of WT1-37 and WT1-126 specific $CD8^+$ T-cell clones was performed as previously described (32). Briefly, peripheral blood mononuclear cells (PBMC) were isolated from blood samples. Subsequently, $CD8^+$ T cells and $CD14^+$ monocytes were isolated from PBMC using magnetic-activated cell sorting. Monocytes were used to generate monocyte-derived DCs. Isolated $CD8^+$ T cells were specifically activated and expanded in two rounds of *in vitro* stimulation (IVS; **Figure 1A (1)**). For the first IVS of 8 days, $CD8^+$ T cells were co-cultured with autologous monocyte-derived DCs pulsed with HLA-A*02:01-restricted WT1-37 or WT1-126 peptide in a 10:1 T cell:DC ratio. For the second IVS, primed $CD8^+$ T cells were co-cultured for 8 days with irradiated autologous WT1-37 or WT1-126 peptide-pulsed CD14 and CD8-depleted peripheral blood lymphocytes (PBL). Both co-cultures were initiated in RPMI with 10% human AB (hAB) serum supplemented with interleukin (IL)-21 (Immunotools, Friesoythe, Germany) (34). Every 2-3 days cells were passaged in RPMI with 10% hAB supplemented with IL-7 and IL-15. After 16 days, cells were harvested and washed for bulk-cell sorting of WT1-37 or WT1-126-reactive T-cell clones (**Figure 1A (2)**). Harvested T cells were stained with anti-human CD3-PerCP-Cy5.5, CD8-Pacific Blue and APC-labeled WT1-37 or WT1-126 HLA-A*02:01 tetramers (kindly provided by Prof. David A. Price). CD14-FITC and CD19-FITC were added to the staining panel to gate out remaining monocytes and B cells. All monoclonal antibodies are purchased from BD Biosciences (Erembodegem, Belgium). Fixable Aqua dead cell stain (ThermoFisher Scientific, Merelbeke, Belgium) was used to discriminate between viable and dead cells. At least 5000 antigen-specific T cells were sorted directly into 100 µL RNA-shield (Zymo Research, Irvine, USA ) using a FACSAria II flow cytometric cell sorter (BD Biosciences, Erembodegem, Belgium) and stored at -20°C for future use. An example of the applied gating strategy for sorting WT1-specific $CD8^+$ T cells is depicted in **Figure S1**.

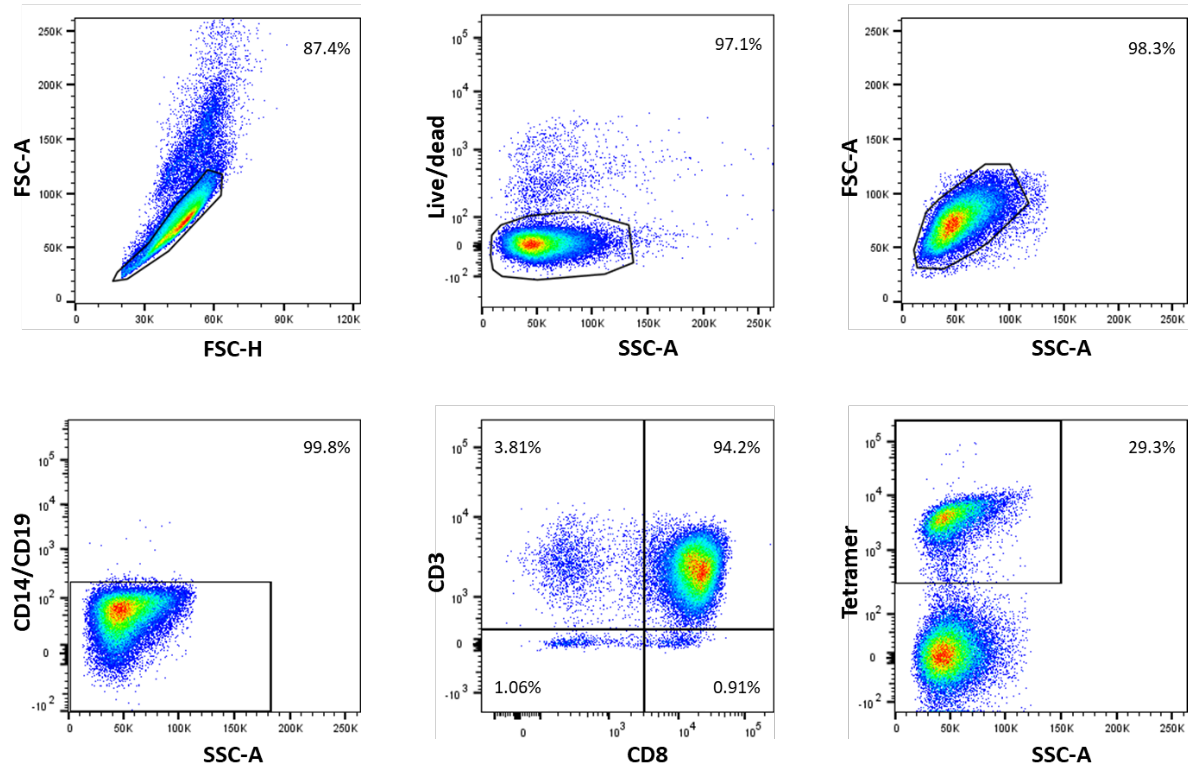


**Figure S1: Gating strategy used for flow cytometric cell sorting of antigen-specific CD8+ T cells.** Plots show the gating strategy used for cell sorting of APC-conjugated tetramer-labeled antigen-specific CD8+ T cells. Viable cells were identified by first gating on the single cells, followed by gating on Fixable Aqua dead cell stain negative population and gating on the forward scatter (FSC) and side scatter (SCC) density plots. An FITC dump channel is used to gate out remaining monocytes and B cells. Subsequently, CD8+ T cells were gated within the viable CD14-CD19- population using a CD3 vs CD8 quadrant plot. Lastly, the population of antigen-specific CD8+ T cells was identified using APC-labeled peptide-specific MHC class I tetramers. The threshold to determine the tetramer positive, and thus antigen-specific CD8+ T cells, was based on a non-stimulated negative control condition.

**TCR sequencing of sorted T cells**

TCR cDNA library preparation and sequencing (**Figure 1A (3)**) was done as previously described (35). RNA was extracted from sorted WT1-37-specific and WT1-126-specific T cells using Quick-RNA Microprep kit (Zymo Research, Irvine, USA). Extracted RNA was immediately used for RNA-based library preparation. The QIAseq Immune Repertoire RNA Library kit (Qiagen, Venlo, The Netherlands) amplifies TCRα, -β, -γ, and -δ chains. After quality control using a Fragment Analyzer (Agilent, Santa Clara, CA), concentration of the cDNA was measured with the Qubit 1x HS DNA Assay kit (Thermo Fisher Scientific, Waltham, MA) and pools were equimolarly pooled and prepared for sequencing on the NextSeq platform (Illumina, San Diego, CA).

**Generation of a WT1-specific database**

Following mini-bulk TCR sequencing, MiXCR (36) (version 3.0.7) was applied to convert the raw reads into TCR sequences. From the resulting MiXCR files, TCR β sequences were collected and parsed similar to our standard TCRex parsing pipeline (28). Parsing steps included the removal of TCRs (i) containing CDR3 beta sequences that were not surrounded by the conserved Cysteine and Phenylalanine residues, (ii) containing a stop codon or another non-amino acid character, and (iii) sequences with orphon genes (37) and removal of the allele info from the V and J genes. In the case of multiple V/J genes, the first gene was selected for every CDR3 beta sequence. For TCRs matched with multiple epitopes, the right epitope partner was identified. To this end, TCR read counts were compared between the associated epitopes. In short, the highest TCR read count was selected for every epitope. In case the fraction between the two most abundant epitope-specific clones was at least 100, the epitope with the highest read count was selected as the true epitope partner. All CDR3 beta sequences with fractions below 100, were removed entirely from the database. The final database is referred to as 'in-house WT1-TCR DB' throughout this paper (**Figure 1A (4)**).

**Training of WT1-specific prediction models**

WT1-specific models were trained using our in-house WT1-TCR DB by the TCRex tool (**Figure 1A (5)**) (28). In brief, TCRex trains a random forest classifier for every epitope separately using a positive training data set (i.e., list of epitope-specific TCRs) and a negative training data set (i.e., a list of TCRs collected from bulk TCR repertoires from healthy donors), the latter by default integrated in the TCRex tool. After model training, each TCRex model predicts for every TCR in a repertoire if it recognizes the specific epitope. The resulting score is compared to the score distribution of a built-in background dataset of 100 000 TCRs to evaluate how many of the background TCRs have a score equal or higher than the TCR of interest. This is summarized as a baseline prediction rate (BPR) value and reported by TCRex for every studied TCR-epitope pair. Hence, a BPR threshold can be applied to retain only those TCR-epitope pairs with a BPR score equal or below this threshold. By default, the threshold is set to 0.01% meaning that for every predicted epitope-specific TCR maximum 10 TCRs in the background data set have a score higher or equal to this predicted epitope-specific TCR. To reduce redundancy in the training data sets due to TCR sequences with identical CDR3 beta sequences but different V/J genes, the TCR sequence with the highest read count was retained for every duplicated CDR3 beta region. The final models were evaluated using the build-in cross-validation strategy and performance metrics of TCRex.

**Evaluation of the publicity of WT1-specific CDR3 beta sequences**

To investigate the level of public TCRs in our in-house WT1-TCR DB, CDR3 beta sequences obtained after two WT1-peptide *in vitro* stimulations that are shared by more than one healthy donor were identified (**Figure 1B (6)**). Here, public TCRs are defined as TCRs having a CDR3 beta sequence occurring in at least two out of twelve and two out of seven healthy donors for WT1-37 and WT1-126, respectively.

**Clustering and TCR motif discovery**

To evaluate TCR similarity of WT1-specific TCRs over the different healthy donors, the TCRs in our in-house WT1-TCR DB were clustered for each epitope according to their CDR3 beta sequence with clusTCR (29) (version 0+untagged.107.g15006f4) and visualized with the spring layout function of NetworkX (version 2.5.1; **Figure 1B (7)**). ClusTCR groups TCRs together based on their CDR3 beta amino acid sequences and a Hamming distance of one, i.e., a maximum of one amino acid difference between any two connected TCRs. By assigning a specific color to every donor and depicting public CDR3s in black, inter-individual clusters were visualized. Amino acid logos were created for the largest clusters.

**Examination of the V/J gene usage of WT1-specific TCR sequences**

To perform an enrichment analysis on the V and J genes of the WT1-specific TCR repertoire, an independent background data set was needed consisting of naïve TCRs sequenced from healthy individuals using a protocol similar to the identification of the WT1-specific TCRs (i.e., RNA-based sequencing and identification with MiXCR (36)). This data was searched for in the iReceptor Gateway (38) on 26$^{th}$ January 2022 with following filter steps: Organism = Homo sapiens; PCR target = TRB; cell subset: CD8-positive, alpha-beta T cell; T cell; effector CD8-positive, alpha-beta T cell; naïve thymus-derived CD8-positive, alpha-beta T cell; Tissue: blood; peripheral blood, venous blood; Target substrate: RNA. After removal of all entries associated with diseases, specific T cells, memory T cells, CD4 T cells or TCRs identified with another tool than MiXCR, only one study remained containing more than one suitable TCR repertoire sample. From this study (39), the samples containing CD8$^{+}$ T cells at day 0 were downloaded. All of these TCR repertoire samples were derived from healthy individuals with at least one HLA-A*02:01 allele, which presents both WT1 epitopes. The final background TCR repertoire was created by removing all non-productive sequences, parsing the CDR3 beta sequences and V/J genes similar to our in-house WT1-TCR DB and removing all duplicate TCRs (i.e., identical CDR3 beta sequences and V/J genes). Enrichment of V/J genes in the WT1-specific TCR repertoire was assessed by comparing the occurrence of every gene for all sequenced, unique TCR sequences (i.e., all unique combinations of CDR3 beta sequences and V/J genes) in our in-house WT1-TCR DB with the background TCR repertoire (**Figure 1B (8)**). For every V/J gene and WT1 epitope, the number of occurrences in the WT1-specific repertoire with its occurrences in the background repertoire was evaluated with Fisher exact tests. To avoid enrichment results for V/J genes with only one count, the enrichment analysis was restricted to V/J genes which were at least two times associated with the studied WT1-epitope in the WT1-specific dataset. Following Benjamini-Hochberg correction on all V genes or J genes per epitope, enriched V/J genes were identified.

**Identification of WT1-specific TCRs in independent cancer repertoires**

All WT1-specific TCRs present in the training data set were derived from healthy donors (in-house WT1-TCR DB). To evaluate whether similar TCRs are present in cancer patients, we searched for WT1-specific TCRs in the repertoires from an independent AML patient study. This independent study was retrieved from the TCRdb (40) after searching for studies in AML sharing clinical response information (i.e., complete remission or relapse). This study aimed to investigate the characteristics of bone marrow T cells in two distinct patient groups: those with relapsing AML and those who achieved complete remission after undergoing hematopoietic stem cell transplantation. (33). TCR β sequence data was available for a subset of the participants included in the study, consisting of three AML patients with relapse, three AML patients who achieved complete remission and three healthy individuals. All TCR data was derived from the bone marrow of these individuals and consisted of a combination of $CD4^+$ and $CD8^+$ T cell-derived TCRs. This data (project ID: PRJNA510967) was downloaded from the TCRdb (40). For each individual, all TCRs were collected and converted into TCRex format. In case multiple samples were present for a single individual, all TCR data was combined and analyzed as a single repertoire. To identify WT1-specific TCRs, a look-up approach was used. Here, TCRs containing a CDR3 beta sequence which exactly matched one of the CDR3 beta sequences in our in-house WT1-TCR DB were considered WT1 specific. Of note, this method does not allow the identification of TCRs having 'unseen' CDR3 beta sequences, i.e., sequences that were not detected in the laboratory and thus are not present in the in-house WT1-TCR DB. To increase the general identification rate of WT1-specific TCRs in the small repertoires, all repertoires were analyzed with the two trained TCRex models and the default BPR threshold of 0.01 %. Finally, all repertoires were clustered with clusTCR (29) to identify those clusters with at least one WT1-specific TCR.

**Statistical analysis**

The enrichment analysis was performed in Python 3.6.10 using the fisher_exact function from the SciPy package (version 1.5.2)(41). R version 3.6.2 was used to perform multiple testing correction. P values < 0.05 after multiple testing correction were considered significant.

#### S2: Overview of additional files

In addition to the relevant code of this project, the github repository stores relevant data, that might be interesting for others.

##### (1) Overview of all training data

The folder data/training_data contains all WT1-specific TCRs that were used to train the models. The sequences for the training of the WT1-37 specific model are stored in the folder WT1-37 while the sequences for the WT1-126 model are stored in the WT1-126 folder.

##### (2) Overview of public TCRs in healthy volunteers

Results/public_cdrs/public_126.tsv and public_37.tsv give an overview of the public CDR3 beta sequences, their V/J genes and their count for every volunteer. CDR3 beta sequences derived from identical RNA sequences are placed within the square brackets.

#### S3: Summary of sequencing and TCR clonotyping

The main text describes the isolation and sequencing of the TCR beta sequence of $WT1_{37\text{-}45}$ and $WT1_{126\text{-}134}$ specific T-cells derived from healthy volunteers. Table S1 gives an overview of the number of raw reads for every volunteer and the resulting number of clonotypes as identified by MiXCR.

**Table S1:** Overview of the number of raw reads and the number of clonotypes identified by MiXCR.

| Epitope | Volunteer | Number of raw reads | Number of TCR clones identified by MiXCR |
|---|---|---|---|
| WT1-126 | DR12 | 1306013 | 309 |
| | DR15 | 592986 | 115 |
| | DR24 | 465023 | 219 |
| | DR25 | 199401 | 80 |
| | DR26 | 52732 | 54 |
| | DR50 | 1929187 | 156 |
| | DR55 | 683538 | 103 |
| WT1-37 | DR10 | 2978042 | 1976 |
| | DR12 | 959882 | 1787 |
| | DR18 | 3205638 | 579 |
| | DR23 | 1500212 | 1222 |
| | DR24 | 1259270 | 791 |
| | DR25 | 1801565 | 1204 |
| | DR26 | 1296209 | 890 |
| | DR47 | 867115 | 113 |
| | DR48 | 464019 | 46 |
| | DR50 | 1359068 | 158 |
| | DR55 | 1323693 | 155 |
| | DR56 | 2919444 | 285 |

#### S4: WT1-specific TCR CDR3 beta sequences are shared across healthy individuals

As shown in **Figure S2**, CDR3 beta sequences of the identified WT1-specific TCRs were frequently shared between donors. When considering identical CDR3 beta sequences (i.e., when V/J genes were not taken into consideration), 234 out of 1262 (18.5%) sequences of the WT1-37 specific repertoire and 27 out of 101 (26.7%) sequences of the WT1-126 specific repertoire were public. More than 90% of these public TCRs were also shared between two or three donors (**Figure S2**). An overview of all identified public TCRs including their V/J genes and read count per donor is provided in **Supplemental material S2**.

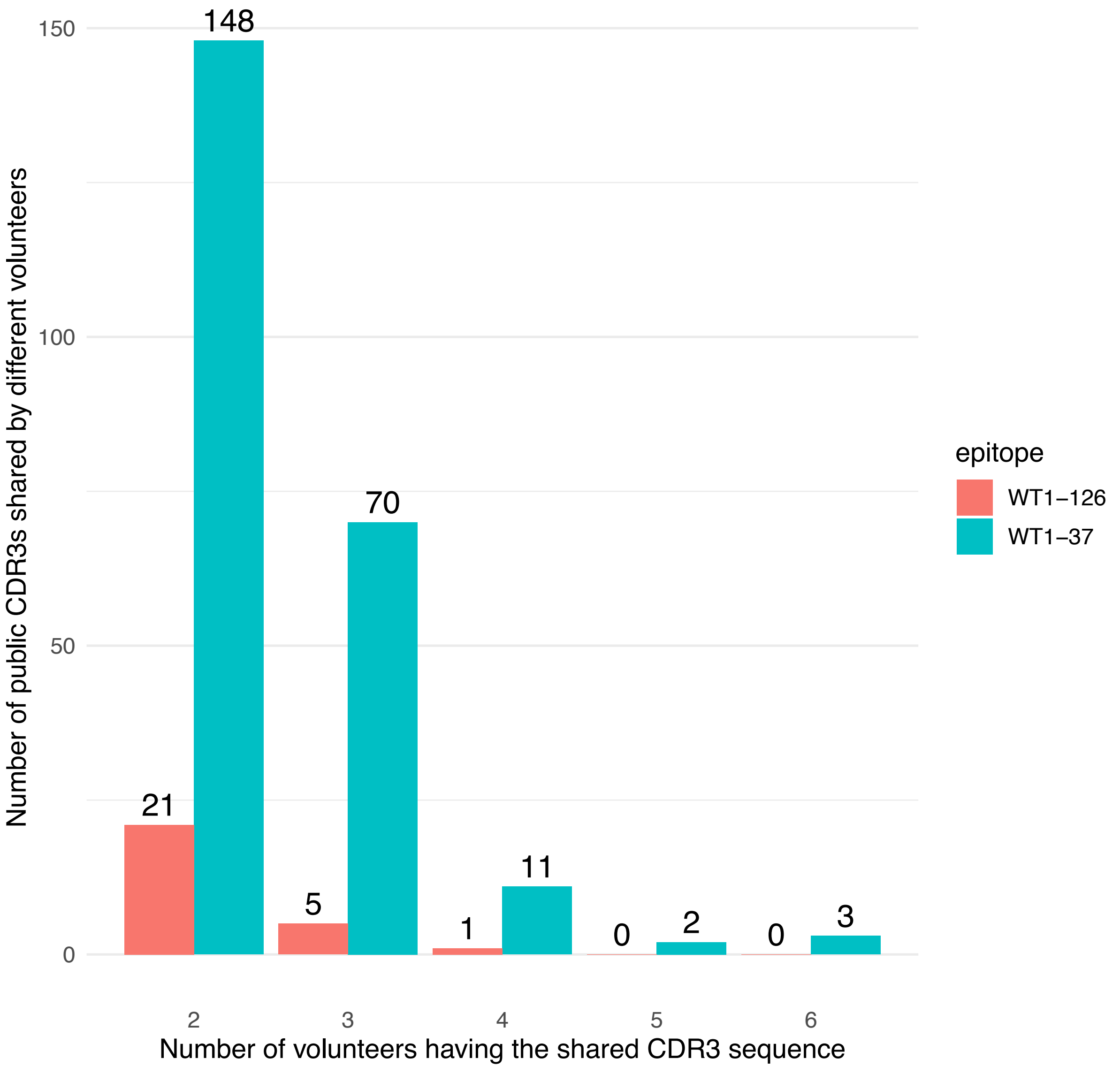


**Figure S2: Number of public WT1-specific TCR CDR3 beta sequences in healthy donors.** Quantification of CDR3 beta sequences of *in vitro* expanded WT1-37 specific (blue bars) and WT1-126 specific (red bars) T cells that are shared between at least two healthy donors (referred to as 'public'). The x-axis represents the number of donors for each shared CDR3 beta sequence.

#### S5: Sequence logos of WT1-specific training data

After clustering the training TCRs for each WT1-epitope specifically, sequence logos were created for the largest clusters, i.e. clusters with at least 3 TCRs for WT1-126 and at least 4 TCRs for WT1-37.

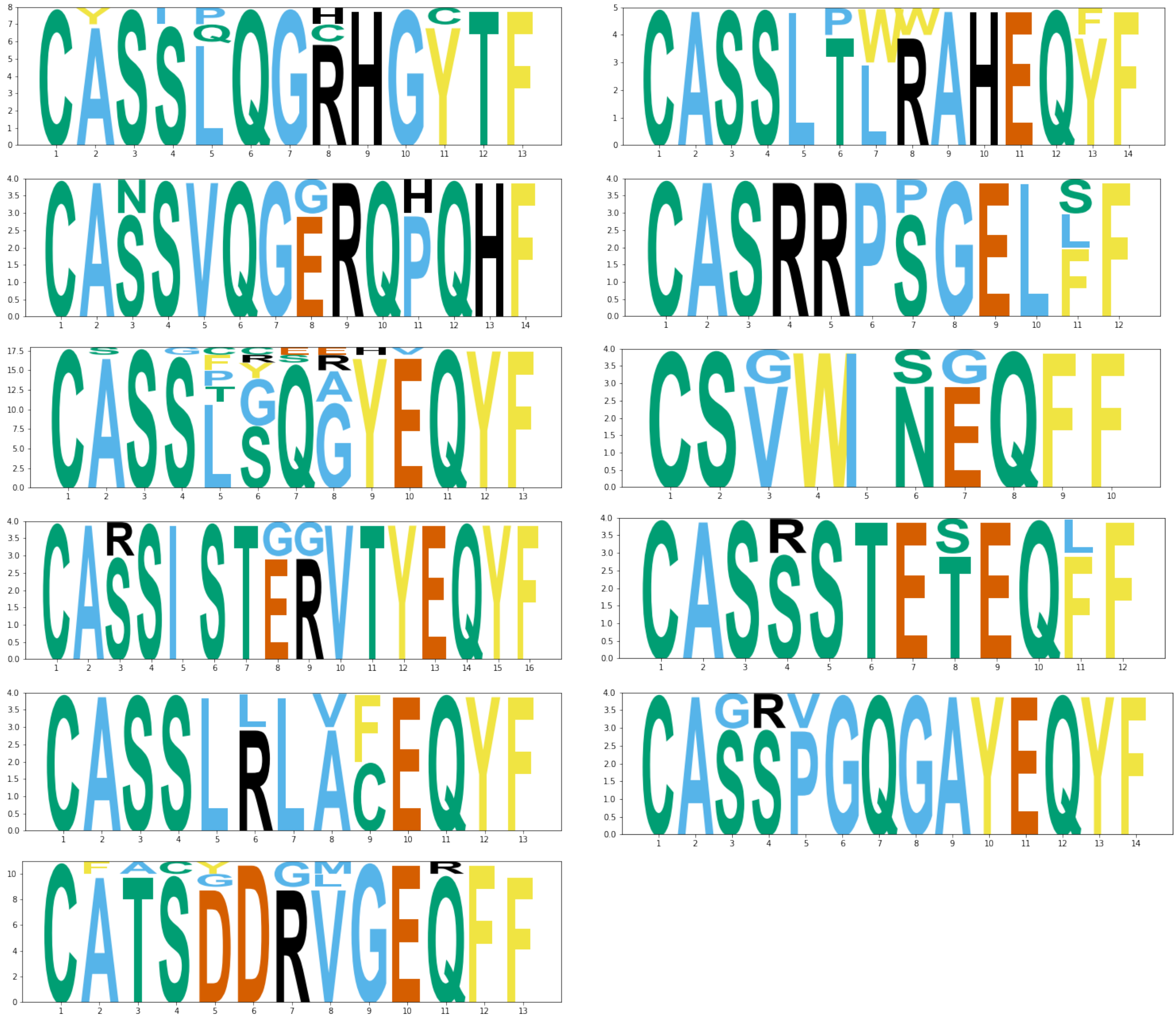

**Figure S3:** Sequence logos for all WT1-37 TCRs clusters with at least 4 TCRs

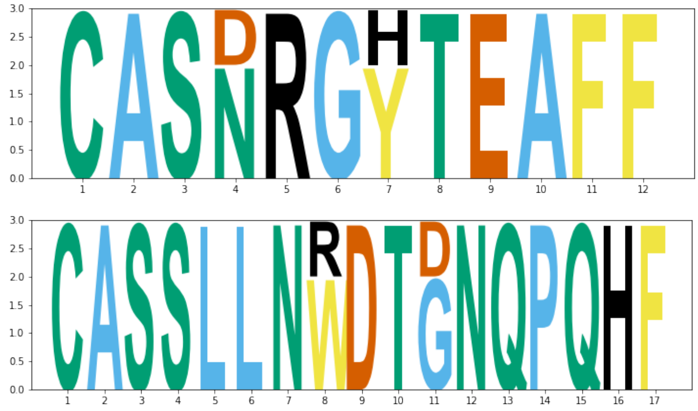

**Figure S4:** Sequence logos for all WT1-126 TCRs clusters with at least 3 TCRs

#### S6: V/J gene distributions differ between WT1-37 and WT1-126 specific TCRs

It has been described for antigens, like Melan-A and MELOE-1, that some V/J genes in antigen-specific TCRs are more prevalent than others (42). To verify the same hypothesis for WT1-specific TCRs, frequencies of every V and J gene present in our in-house WT1-TCR DB were quantified (**Figure S5**). As seen in **Figure S5A-5D**, there were noticeable differences in the distributions and thus the frequencies of the different V/J genes and families. To check for possible enrichment of the various V/J genes and families, the occurrence of each gene/family was compared with the occurrence of a representative background repertoire consisting of naïve TCRs from healthy individuals (39). In short, a Fisher's exact test was performed for every V/J gene or family and WT1-epitope where the number of occurrences in the WT1-specific repertoire was compared with the occurrences in the background repertoire. The Benjamini-Hochberg corrected p-values are shown in **Supplemental material S7**. This enrichment analysis revealed a significant elevated level of TRBJ02-07 and 12 TRBV genes (TRBV19, TRBV07-03, TRBV28, TRBV27, TRBV06-03, TRBV24-01, TRBV30, TRBV02, TRBV05-07, TRBV12-01, TRBV18, TRBV06-07) for the WT1-37 specific TCRs, and 4 TRBV genes (TRBV05-04, TRBV05-08, TRBV07-09, TRBV30) for the WT1-126 specific TCRs (**Supplemental material S7**). One J gene, TRBJ02-07, occurred more often within WT1-37 specific TCRs. These data confirm that a number of V/J genes are more often present specifically in WT1-specific TCRs.

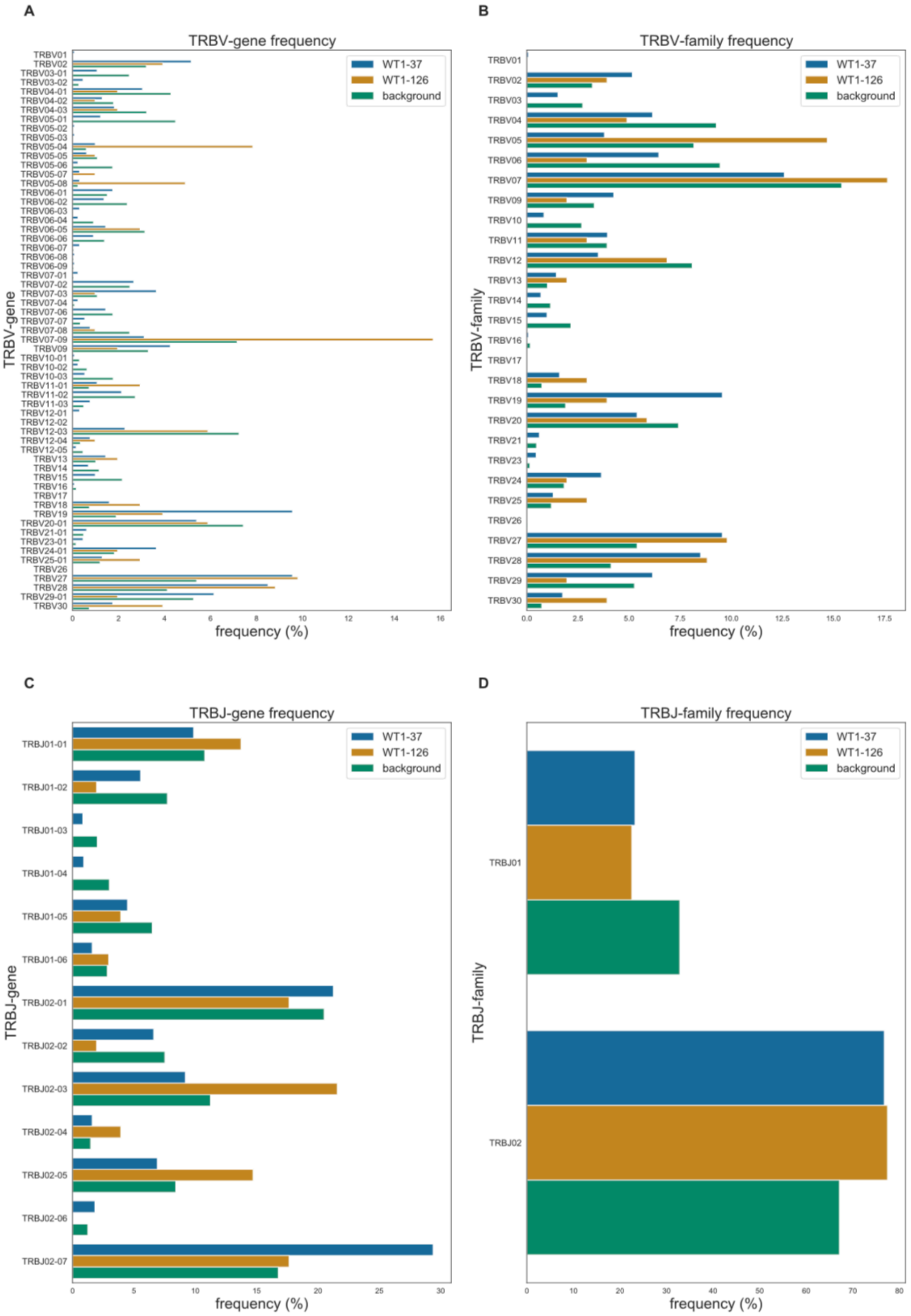


**Figure S5: Comparison of V/J gene (A, C) and V/J family (B, D) distribution with representative background TCR repertoire.** Bar charts (**A**) and (**C**) show the frequencies of specific TCR β V/J genes, following the international ImMunoGeneTics information system (IMGT) gene nomenclature, for the TCRs derived from healthy donors specific for WT1-37 (blue bars) or WT1-126 (orange bars), while graphs (**B**) and (**D**) show frequencies of TCR β V/J gene families (i.e., IMGT subgroups). To enable the identification of V/J genes that are overrepresented in one of the WT1-specific repertoires, their frequencies were compared with the frequency distribution of all V/J genes for a representative background dataset (green bars). This enables the distinction between gene enrichment or depletion when comparing the gene frequencies between the two epitopes.

#### S7: V/J gene enrichment in WT1-specific TCRs

To study the relative abundance of the V/J genes in the WT1-specific repertoire, the presence of the V/J genes in the WT1-37 and WT1-126 repertoire was compared with a representative background dataset using a Fisher exact test. The background repertoire contained a total of 2 526 973 unique combinations of CDR3 beta and V/J genes, while the WT1-37 and WT1-126 contained respectively 1317 and 102 unique combinations. The adjusted p values (Benjamini-Hochberg correction) are shown in the following 4 tables together with the counts and percentages of every V/J gene in the studied WT1-specific and background dataset. The significant results are marked in green. The p values are rounded up to 3 significant numbers, while all percentages are rounded to two digits after the comma.

**Table S2:** Enrichment results for WT1-37 J genes

| TRBJ gene | WT1-37 counts | WT1-37 percentage | Background counts | Background percentage | Adjusted p value |
|---|---|---|---|---|---|
| TRBJ02-07 | 387 | 29,38 | 423734 | 16,77 | 3,12E-19 |
| TRBJ02-06 | 24 | 1,82 | 31514 | 1,25 | 0,313 |
| TRBJ02-01 | 280 | 21,26 | 518087 | 20,50 | 1 |
| TRBJ02-04 | 21 | 1,59 | 37210 | 1,47 | 1 |
| TRBJ01-01 | 130 | 9,87 | 272116 | 10,77 | 1 |
| TRBJ02-02 | 87 | 6,61 | 190070 | 7,52 | 1 |
| TRBJ02-05 | 91 | 6,91 | 212520 | 8,41 | 1 |
| TRBJ02-03 | 121 | 9,19 | 283983 | 11,24 | 1 |
| TRBJ01-02 | 73 | 5,54 | 195204 | 7,72 | 1 |
| TRBJ01-06 | 21 | 1,59 | 71582 | 2,83 | 1 |
| TRBJ01-05 | 59 | 4,48 | 164188 | 6,50 | 1 |
| TRBJ01-03 | 11 | 0,84 | 51093 | 2,02 | 1 |
| TRBJ01-04 | 12 | 0,91 | 75672 | 2,99 | 1 |

**Table S3:** Enrichment results for WT1-126 J genes

| TRBJ_gene | WT1-126 counts | WT1-126 percentage | Background counts | Background percentage | Adjusted p value |
|---|---|---|---|---|---|
| TRBJ02-03 | 22 | 21,57 | 283983 | 11,24 | 0,0640 |
| TRBJ02-05 | 15 | 14,71 | 212520 | 8,41 | 0,187 |
| TRBJ02-04 | 4 | 3,92 | 37210 | 1,47 | 0,230 |
| TRBJ01-01 | 14 | 13,73 | 272116 | 10,77 | 0,590 |
| TRBJ02-07 | 18 | 17,65 | 423734 | 16,77 | 0,920 |
| TRBJ01-06 | 3 | 2,94 | 71582 | 2,83 | 0,925 |
| TRBJ02-01 | 18 | 17,65 | 518087 | 20,50 | 0,996 |
| TRBJ01-05 | 4 | 3,92 | 164188 | 6,50 | 0,996 |
| TRBJ02-02 | 2 | 1,96 | 190070 | 7,52 | 0,996 |
| TRBJ01-02 | 2 | 1,96 | 195204 | 7,72 | 0,996 |
| TRBJ01-03 | 0 | 0 | 51093 | 2,02 | NA |
| TRBJ01-04 | 0 | 0 | 75672 | 2,99 | NA |
| TRBJ02-06 | 0 | 0 | 31514 | 1,25 | NA |

**Table S4:** Enrichment results for WT1-37 V genes

| TRBV gene | WT1-37 counts | WT1-37 percentage | Background counts | Background percentage | Adjusted p value |
|---|---|---|---|---|---|
| TRBV19 | 126 | 9,57 | 47959 | 1,90 | 2,39E-43 |
| TRBV07-03 | 48 | 3,64 | 27045 | 1,07 | 4,96E-11 |
| TRBV28 | 112 | 8,50 | 104124 | 4,12 | 2,89E-10 |
| TRBV27 | 126 | 9,57 | 136330 | 5,39 | 1,03E-07 |
| TRBV06-03 | 4 | 0,30 | 219 | 0,01 | 6,93E-05 |
| TRBV24-01 | 48 | 3,64 | 46074 | 1,82 | 0,000114 |
| TRBV30 | 23 | 1,75 | 18371 | 0,73 | 0,00128 |
| TRBV02 | 68 | 5,16 | 81012 | 3,21 | 0,00133 |
| TRBV05-07 | 4 | 0,30 | 570 | 0,02 | 0,00148 |
| TRBV12-01 | 4 | 0,30 | 692 | 0,03 | 0,00275 |
| TRBV18 | 21 | 1,59 | 18597 | 0,74 | 0,00539 |
| TRBV06-07 | 4 | 0,30 | 1094 | 0,04 | 0,0121 |
| TRBV07-01 | 3 | 0,23 | 950 | 0,04 | 0,0543 |
| TRBV23-01 | 6 | 0,46 | 3737 | 0,15 | 0,0543 |
| TRBV12-04 | 10 | 0,76 | 8676 | 0,34 | 0,0609 |
| TRBV09 | 56 | 4,25 | 83323 | 3,30 | 0,129 |
| TRBV05-04 | 13 | 0,99 | 15118 | 0,60 | 0,177 |
| TRBV13 | 19 | 1,44 | 25521 | 1,01 | 0,238 |
| TRBV07-04 | 3 | 0,23 | 2036 | 0,08 | 0,241 |
| TRBV11-01 | 14 | 1,06 | 18081 | 0,72 | 0,241 |
| TRBV29-01 | 81 | 6,15 | 133052 | 5,27 | 0,241 |
| TRBV11-03 | 10 | 0,76 | 12290 | 0,49 | 0,268 |
| TRBV03-02 | 6 | 0,46 | 6744 | 0,27 | 0,322 |
| TRBV07-07 | 7 | 0,53 | 8563 | 0,34 | 0,351 |
| TRBV06-01 | 23 | 1,75 | 38172 | 1,51 | 0,558 |
| TRBV21-01 | 8 | 0,61 | 12017 | 0,48 | 0,577 |
| TRBV05-08 | 4 | 0,30 | 5774 | 0,23 | 0,671 |
| TRBV07-02 | 35 | 2,66 | 63056 | 2,50 | 0,692 |
| TRBV25-01 | 17 | 1,29 | 30246 | 1,20 | 0,724 |
| TRBV07-06 | 19 | 1,44 | 44287 | 1,75 | 1 |
| TRBV11-02 | 28 | 2,13 | 68812 | 2,72 | 1 |
| TRBV04-02 | 17 | 1,29 | 44959 | 1,78 | 1 |
| TRBV06-06 | 12 | 0,91 | 34978 | 1,38 | 1 |
| TRBV14 | 9 | 0,68 | 29196 | 1,16 | 1 |
| TRBV05-05 | 8 | 0,61 | 27312 | 1,08 | 1 |
| TRBV12-05 | 2 | 0,15 | 11463 | 0,45 | 1 |
| TRBV10-02 | 3 | 0,23 | 15873 | 0,63 | 1 |
| TRBV04-01 | 40 | 3,04 | 108058 | 4,28 | 1 |

| | | | | | |
|---|---|---|---|---|---|
| TRBV06-02 | 18 | 1,37 | 60039 | 2,38 | 1 |
| TRBV20-01 | 71 | 5,39 | 187630 | 7,43 | 1 |
| TRBV04-03 | 24 | 1,82 | 81383 | 3,22 | 1 |
| TRBV06-04 | 3 | 0,23 | 23058 | 0,91 | 1 |
| TRBV15 | 13 | 0,99 | 54529 | 2,16 | 1 |
| TRBV03-01 | 14 | 1,06 | 62312 | 2,47 | 1 |
| TRBV06-05 | 19 | 1,44 | 79413 | 3,14 | 1 |
| TRBV10-03 | 7 | 0,53 | 44588 | 1,76 | 1 |
| TRBV07-08 | 10 | 0,76 | 62651 | 2,48 | 1 |
| TRBV05-06 | 3 | 0,23 | 43914 | 1,74 | 1 |
| TRBV07-09 | 41 | 3,11 | 180882 | 7,16 | 1 |
| TRBV05-01 | 16 | 1,21 | 113111 | 4,48 | 1 |
| TRBV12-03 | 30 | 2,28 | 183014 | 7,24 | 1 |
| TRBV01 | 1 | 0,08 | 774 | 0,03 | NA |
| TRBV05-02 | 1 | 0,08 | 0 | 0,00 | NA |
| TRBV05-03 | 1 | 0,08 | 763 | 0,03 | NA |
| TRBV06-08 | 1 | 0,08 | 940 | 0,04 | NA |
| TRBV06-09 | 1 | 0,08 | 1068 | 0,04 | NA |
| TRBV10-01 | 1 | 0,08 | 7453 | 0,29 | NA |
| TRBV16 | 1 | 0,08 | 4097 | 0,16 | NA |
| TRBV12-02 | 0 | 0,00 | 713 | 0,03 | NA |
| TRBV17 | 0 | 0,00 | 57 | 0,00 | NA |
| TRBV26 | 0 | 0,00 | 203 | 0,01 | NA |

**Table S5:** Enrichment results for WT1-126 V genes

| TRBV gene | WT1-126 counts | WT1-126 percentage | Background counts | Background percentage | Adjusted p value |
|---|---|---|---|---|---|
| TRBV05-04 | 8 | 7,84 | 15118 | 0,60 | 7,50E-06 |
| TRBV05-08 | 5 | 4,90 | 5774 | 0,23 | 5,41E-05 |
| TRBV07-09 | 16 | 15,69 | 180882 | 7,16 | 0,0359 |
| TRBV30 | 4 | 3,92 | 18371 | 0,73 | 0,0376 |
| TRBV28 | 9 | 8,82 | 104124 | 4,12 | 0,121 |
| TRBV11-01 | 3 | 2,94 | 18081 | 0,72 | 0,121 |
| TRBV18 | 3 | 2,94 | 18597 | 0,74 | 0,121 |
| TRBV27 | 10 | 9,80 | 136330 | 5,39 | 0,154 |
| TRBV25-01 | 3 | 2,94 | 30246 | 1,20 | 0,273 |
| TRBV19 | 4 | 3,92 | 47959 | 1,90 | 0,273 |
| TRBV13 | 2 | 1,96 | 25521 | 1,01 | 0,507 |
| TRBV02 | 4 | 3,92 | 81012 | 3,21 | 0,698 |
| TRBV24-01 | 2 | 1,96 | 46074 | 1,82 | 0,858 |
| TRBV06-05 | 3 | 2,94 | 79413 | 3,14 | 0,891 |
| TRBV12-03 | 6 | 5,88 | 183014 | 7,24 | 0,943 |
| TRBV20-01 | 6 | 5,88 | 187630 | 7,43 | 0,943 |
| TRBV04-03 | 2 | 1,96 | 81383 | 3,22 | 0,943 |
| TRBV09 | 2 | 1,96 | 83323 | 3,30 | 0,943 |
| TRBV04-01 | 2 | 1,96 | 108058 | 4,28 | 0,969 |
| TRBV29-01 | 2 | 1,96 | 133052 | 5,27 | 0,969 |
| TRBV04-02 | 1 | 0,98 | 44959 | 1,78 | NA |
| TRBV05-05 | 1 | 0,98 | 27312 | 1,08 | NA |
| TRBV05-07 | 1 | 0,98 | 570 | 0,02 | NA |
| TRBV07-03 | 1 | 0,98 | 27045 | 1,07 | NA |
| TRBV07-08 | 1 | 0,98 | 62651 | 2,48 | NA |
| TRBV12-04 | 1 | 0,98 | 8676 | 0,34 | NA |
| TRBV01 | 0 | 0 | 774 | 0,03 | NA |
| TRBV03-01 | 0 | 0 | 62312 | 2,47 | NA |
| TRBV03-02 | 0 | 0 | 6744 | 0,27 | NA |
| TRBV05-01 | 0 | 0 | 113111 | 4,48 | NA |
| TRBV05-03 | 0 | 0 | 763 | 0,03 | NA |
| TRBV05-06 | 0 | 0 | 43914 | 1,74 | NA |
| TRBV06-01 | 0 | 0 | 38172 | 1,51 | NA |
| TRBV06-02 | 0 | 0 | 60039 | 2,38 | NA |
| TRBV06-03 | 0 | 0 | 219 | 0,01 | NA |
| TRBV06-04 | 0 | 0 | 23058 | 0,91 | NA |
| TRBV06-06 | 0 | 0 | 34978 | 1,38 | NA |
| TRBV06-07 | 0 | 0 | 1094 | 0,04 | NA |
| TRBV06-08 | 0 | 0 | 940 | 0,04 | NA |
| TRBV06-09 | 0 | 0 | 1068 | 0,04 | NA |

| | | | | | |
|---|---|---|---|---|---|
| TRBV07-01 | 0 | 0 | 950 | 0,04 | NA |
| TRBV07-02 | 0 | 0 | 63056 | 2,50 | NA |
| TRBV07-04 | 0 | 0 | 2036 | 0,08 | NA |
| TRBV07-06 | 0 | 0 | 44287 | 1,75 | NA |
| TRBV07-07 | 0 | 0 | 8563 | 0,34 | NA |
| TRBV10-01 | 0 | 0 | 7453 | 0,29 | NA |
| TRBV10-02 | 0 | 0 | 15873 | 0,63 | NA |
| TRBV10-03 | 0 | 0 | 44588 | 1,76 | NA |
| TRBV11-02 | 0 | 0 | 68812 | 2,72 | NA |
| TRBV11-03 | 0 | 0 | 12290 | 0,49 | NA |
| TRBV12-01 | 0 | 0 | 692 | 0,03 | NA |
| TRBV12-02 | 0 | 0 | 713 | 0,03 | NA |
| TRBV12-05 | 0 | 0 | 11463 | 0,45 | NA |
| TRBV14 | 0 | 0 | 29196 | 1,16 | NA |
| TRBV15 | 0 | 0 | 54529 | 2,16 | NA |
| TRBV16 | 0 | 0 | 4097 | 0,16 | NA |
| TRBV17 | 0 | 0 | 57 | 0,00 | NA |
| TRBV21-01 | 0 | 0 | 12017 | 0,48 | NA |
| TRBV23-01 | 0 | 0 | 3737 | 0,15 | NA |
| TRBV26 | 0 | 0 | 203 | 0,01 | NA |

#### S8: Overview of the performance metrics of the trained TCRex models

Table S6 shows the performance metrics for the two WT1 models trained by TCRex.

**Table S6:** Overview of the performance metrics of the trained TCRex models

| Epitope | Size of TCR dataset | Balanced accuracy | AUROC | Average precision |
|---|---|---|---|---|
| WT1-37 | 1262 | 0.56 ± 0.01 | 0.8 ± 0.02 | 0.49 ± 0.03 |
| WT1-126 | 101 | 0.58 ± 0.03 | 0.69 ± 0.11 | 0.49 ± 0.15 |

Rows show mean ± SD for every performance metric. Abbreviations: AUROC, area under the ROC curve; TCR, T cell receptor; WT1, Wilms' tumor protein 1.

#### S9: Identified WT1-specific TCR sequences in the AML study

The following tables lists the matched training TCR sequences (table S7) in the TCR repertoires of every volunteer from the AML study and the predicted WT1-specific TCRs (tables S8). It shows that some CDR3 beta sequences occur in multiple volunteers with other V genes (colored green or yellow). In addition, 2 TCR sequences were identified both with the look up method and the TCRex models (TRBV28-CASSPGYEQYF-TRBJ02-07 and TRBV06-03-CASSLGTEAFF- TRBJ01-01)

**Table S7:** WT1-specific TCR sequences identified using the look-up method for AML patients and healthy volunteers from the described AML study. To evaluate whether the TCRs with CDR3 beta sequences identical to our in-house WT1-TCR DB also contained the same V/J genes as these database TCRs, the genes for the TCR and its database match are shown in the table (i.e. 'x' denotes the features in the volunteers of the AML study , while 'y' denotes the features of the sequences in the in-house WT1-TCR DB )

| TRBV gene x | CDR3 beta | TRBJ gene x | epitope | TRBV gene y | TRBJ gene y | Volunteer |
|---|---|---|---|---|---|---|
| TRBV27 | CASSLGGNQPQHF | TRBJ01-05 | WT1-37 | TRBV27 | TRBJ1-5 | PT1_CR1 |
| TRBV07-03 | CASSPQDGYEQYF | TRBJ02-07 | WT1-37 | TRBV19 | TRBJ2-7 | PT1_CR1 |
| TRBV05-06 | CASSLQGYSNQPQHF | TRBJ01-05 | WT1-37 | TRBV7-2 | TRBJ1-5 | PT1_CR1 |
| TRBV06-05 | CASSSGTGAYEQYF | TRBJ02-07 | WT1-37 | TRBV19 | TRBJ2-7 | PT1_CR1 |
| TRBV07-02 | CASSLGGNQPQHF | TRBJ01-05 | WT1-37 | TRBV27 | TRBJ1-5 | PT5_REL2 |
| TRBV05-08 | CASSLQGYSNQPQHF | TRBJ01-05 | WT1-37 | TRBV7-2 | TRBJ1-5 | PT5_REL2 |
| TRBV07-06 | CASSLGGNQPQHF | TRBJ01-05 | WT1-37 | TRBV27 | TRBJ1-5 | PT6_REL3 |
| TRBV05-01 | CASPFPLCSYNEQFF | TRBJ02-01 | WT1-37 | TRBV13 | TRBJ2-1 | PT3_CR3 |
| TRBV28 | CASSPGYEQYF | TRBJ02-07 | WT1-37 | TRBV28 | TRBJ2-7 | PT2_CR2 |
| TRBV05-01 | CASSPGQGYEQYF | TRBJ02-07 | WT1-37 | TRBV7-6 | TRBJ2-7 | PT2_CR2 |
| TRBV06-06 | CASSLGSNQPQHF | TRBJ01-05 | WT1-37 | TRBV7-9 | TRBJ1-5 | PT2_CR2 |
| TRBV06-03 | CASSLGTEAFF | TRBJ01-01 | WT1-126 | TRBV27 | TRBJ1-1 | PT2_CR2 |
| TRBV12-04 | CASRPGQGAYEQYF | TRBJ02-07 | WT1-37 | TRBV19 | TRBJ2-7 | HD3 |

**Table S8:** WT1-specific TCR sequences identified using the TCRex prediction models for AML patients and healthy volunteers from the described AML study

| TRBV gene | CDR3 beta | TRBJ gene | epitope | Volunteer |
|---|---|---|---|---|
| TRBV07-08 | CASSLGQAYEQYF | TRBJ02-07 | WT1-37 | PT6_REL3 |
| TRBV07-09 | CASSLLAGEQETQYF | TRBJ02-05 | WT1-126 | PT6_REL3 |
| TRBV28 | CASSPGYEQYF | TRBJ02-07 | WT1-37 | PT2_CR2 |
| TRBV06-03 | CASSLGTEAFF | TRBJ01-01 | WT1-126 | PT2_CR2 |
| TRBV29-01 | CSVEGGSSYEQYF | TRBJ02-07 | WT1-37 | HD3 |
| TRBV24-01 | CATSELAGDVETQYF | TRBJ02-05 | WT1-126 | HD3 |

To see whether the identified TCRs were already identified previously with other epitope partners, they were searched through the VDJdb. To this end, all human TRB sequences (both MHCI and MHCII) were downloaded from the VDJdb site at 09/03/2023. The results are shown in table S9.

**Table S9:** Overview of WT1-specific TCR sequences from tables S7 and S8 that are also represented in the VDJdb. The first column shows the WT1-specific CDR3 beta sequences identified in the AML study that were also found in the VDJdb. For every of these TCRs, their respective V/J genes and WT1-epitope is shown. For the same CDR3 beta sequences, VDJdb information about the HLA background, the V/J genes and its epitope partner is given.

| | Identified WT1-specific TCRs | | VDJdb entries | | | | |
|---|---|---|---|---|---|---|---|
| **CDR3 beta** | **Epitope** | **TRBV/J genes** | **MHC A** | **Epitope** | **Epitope gene** | **Epitope species** | **TRBV/J genes** |
| CASSLGGNQPQHF | WT1-37 | TRBV27_TRBJ01-05, TRBV07-02_TRBJ01-05, TRBV07-06_TRBJ01-05 | HLA-A*01:01 | LTDEMIAQY | Spike | SARS-CoV-2 | TRBV12-3_TRBJ1-5 |
| CASSLGQAYEQYF | WT1-37 | TRBV07-08_TRBJ02-07 | HLA-B*08:01:29 | FLRGRAYGL | EBNA3A | EBV | TRBV7-8_TRBJ2-7 |
| CASSLGQAYEQYF | WT1-37 | TRBV07-08_TRBJ02-07 | HLA-B*44:05:01 | EEYLKAWTF | MLANA | HomoSapiens | TRBV7-8_TRBJ2-7 |
| CASSLGQAYEQYF | WT1-37 | TRBV07-08_TRBJ02-07 | HLA-B*44:05:01 | EEYLQAFTY | ABCD3 | HomoSapiens | TRBV7-8_TRBJ2-7 |
| CASSLGQAYEQYF | WT1-37 | TRBV07-08_TRBJ02-07 | HLA-A*02:01 | GLCTLVAML | BMLF1 | EBV | TRBV7-8_TRBJ2-7 |
| CASSLGQAYEQYF | WT1-37 | TRBV07-08_TRBJ02-07 | HLA-A*01:01 | TTDPSFLGRY | NSP3 | SARS-CoV-2 | TRBV7-8_TRBJ2-7 |
| CASSLGTEAFF | WT1-126 | TRBV06-03_TRBJ01-01 | HLA-A*02 | NLVPMVATV | pp65 | CMV | TRBV5-6_TRBJ1-1 |
| CASSPGQGYEQYF | WT1-37 | TRBV05-01_TRBJ02-07 | HLA-A*02 | NLVPMVATV | pp65 | CMV | TRBV14_TRBJ2-7 |
| CASSPGQGYEQYF | WT1-37 | TRBV05-01_TRBJ02-07 | HLA-A*02:01 | ELAGIGILTV | MLANA | HomoSapiens | TRBV6-3_TRBJ2-7 |
| CASSPGQGYEQYF | WT1-37 | TRBV05-01_TRBJ02-07 | HLA-A*24:02 | NYNYLYRLF | Spike | SARS-CoV-2 | TRBV6-4_TRBJ2-7 |
| CASSPGQGYEQYF | WT1-37 | TRBV05-01_TRBJ02-07 | HLA-A*24:02 | NYNYLYRLF | Spike | SARS-CoV-2 | TRBV5-4_TRBJ2-7 |
| CASSPGYEQYF | WT1-37 | TRBV28_TRBJ02-07 | HLA-A*02:01 | GLCTLVAML | BMLF1 | EBV | TRBV4-1_TRBJ2-7 |
| CASSSGTGAYEQYF | WT1-37 | TRBV06-05_TRBJ02-07 | HLA-A*03:01 | KLGGALQAK | IE1 | CMV | TRBV7-9_TRBJ2-7 |

#### S10: Overview of the CDR3 beta content of clusters containing one or more identified WT1-specific TCRs.

Table S10 lists 7 clusters containing at least one predicted WT1-specific TCR. In table S1, the CDR3 beta content of these 7 clusters is shown, together with the response category of the individual the TCR was sequenced from.

**Table S10:** Overview of the number of CDR3 beta sequences in the clusters containing one or more identified WT1-specific TCRs.

| Cluster | Number of CDR3 beta sequences in every cluster | | | | |
|---|---|---|---|---|---|
| | Healthy | Complete remission | Relapse | WT1-37 TCRs | WT1-126 TCRs |
| 1 | 1 | 4 | 13 | 1 | 0 |
| 2 | 0 | 4 | 0 | 0 | 1 |
| 3 | 0 | 3 | 0 | 1 | 0 |
| 4 | 1 | 6 | 0 | 1 | 0 |
| 5 | 0 | 4 | 2 | 1 | 0 |
| 6 | 1 | 1 | 0 | 1 | 0 |
| 7 | 0 | 2 | 1 | 2 | 0 |

**Table S11:** Overview of all CDR3 sequences of the WT1-specific clusters. 1 represents predicted recognition of the defined epitope, while 0 represents no predicted recognition. The 3 green CDR3 beta sequences are identical.

| CDR3 beta | cluster | WT1-37 | WT1-126 | Volunteer | Response |
|---|---|---|---|---|---|
| CASSPGQEQYF | A | 0 | 0 | PT4_REL1 | Relapse |
| CAISEGNEQYF | A | 0 | 0 | PT4_REL1 | Relapse |
| CASGQGNEQFF | A | 0 | 0 | PT1_CR1 | Complete_remission |
| CASSLGQESYF | A | 0 | 0 | PT4_REL1 | Relapse |
| CATSLGNEQYF | A | 0 | 0 | HD2 | Healthy |
| CAISEGNEQFF | A | 0 | 0 | PT4_REL1 | Relapse |
| CASGLGNEQFF | A | 0 | 0 | PT2_CR2 | Complete_remission |
| CATSLGNEQFF | A | 0 | 0 | PT4_REL1 | Relapse |
| CASSIGQEQYF | A | 0 | 0 | PT4_REL1 | Relapse |
| CATRLGNEQFF | A | 0 | 0 | PT1_CR1 | Complete_remission |
| CASSLGQEQYF | A | 0 | 0 | PT4_REL1 | Relapse |
| CAISVGNEQFF | A | 0 | 0 | PT4_REL1 | Relapse |
| CASSLGQEQFF | A | 0 | 0 | PT4_REL1 | Relapse |
| CASSLGNEQFF | A | 0 | 0 | PT5_REL2 | Relapse |
| CAISSGNEQFF | A | 0 | 0 | PT4_REL1 | Relapse |
| CAISLGNEQFF | A | 0 | 0 | PT4_REL1 | Relapse |
| CASSPGYEQYF | A | 1 | 0 | PT2_CR2 | Complete_remission |
| CASSVGQEQYF | A | 0 | 0 | PT4_REL1 | Relapse |
| CASSEGQGYEQYF | B | 0 | 0 | PT2_CR2 | Complete_remission |
| CASSPGQGYEQYF | B | 1 | 0 | PT2_CR2 | Complete_remission |
| CASSPGTSYEQYF | B | 0 | 0 | PT1_CR1 | Complete_remission |

| | | | | | |
|---|---|---|---|---|---|
| CASSPGPGYEQYF | B | 0 | 0 | HD3 | Healthy |
| CASSPGTPYEQYF | B | 0 | 0 | PT1_CR1 | Complete_remission |
| CASSPGTGYEQYF | B | 0 | 0 | PT2_CR2 | Complete_remission |
| CASSPGTGQEQYF | B | 0 | 0 | PT1_CR1 | Complete_remission |
| CASSLYTEAFF | C | 0 | 0 | PT2_CR2 | Complete_remission |
| CASSLGTEAFF | C | 0 | 1 | PT2_CR2 | Complete_remission |
| CASSFGTEAFF | C | 0 | 0 | PT3_CR3 | Complete_remission |
| CASSGYTEAFF | C | 0 | 0 | PT2_CR2 | Complete_remission |
| CASSFGGAYEQYF | D | 0 | 0 | PT2_CR2 | Complete_remission |
| CASSFGGIYEQYF | D | 0 | 0 | PT1_CR1 | Complete_remission |
| CASSLGQAYEQYF | D | 1 | 0 | PT6_REL3 | Relapse |
| CASSFGQTYEQYF | D | 0 | 0 | PT1_CR1 | Complete_remission |
| CASSFGQAYEQYF | D | 0 | 0 | PT4_REL1 | Relapse |
| CASSLGQHYEQYF | D | 0 | 0 | PT1_CR1 | Complete_remission |
| CASSSGTGAPEQYF | E | 0 | 0 | PT1_CR1 | Complete_remission |
| CASSVGTGAYEQYF | E | 0 | 0 | PT2_CR2 | Complete_remission |
| CASSSGTGAYEQYF | E | 1 | 0 | PT1_CR1 | Complete_remission |
| CASSLGGNQPQHF | F | 1 | 0 | PT1_CR1 | Complete_remission, Relapse, Relapse |
| CASSLGGNQPQHF | F | 1 | 0 | PT5_REL2 | Complete_remission, Relapse, Relapse |
| CASSLGGNQPQHF | F | 1 | 0 | PT6_REL3 | Complete_remission, Relapse, Relapse |
| CASSLGSNQPQHF | F | 1 | 0 | PT2_CR2 | Complete_remission |
| CSVEGGSSYEQYF | G | 1 | 0 | HD3 | Healthy |
| CSVEGLSSYEQYF | G | 0 | 0 | PT1_CR1 | Complete_remission |
